# Numerical and Spatial Patterning of Yeast Meiotic DNA Breaks by Tel1

**DOI:** 10.1101/059022

**Authors:** Neeman Mohibullah, Scott Keeney

## Abstract

The Spo11-generated double-strand breaks (DSBs) that initiate meiotic recombination are dangerous lesions that can disrupt genome integrity, so meiotic cells regulate their number, timing, and distribution. Here, we use Spo11-oligonucleotide complexes, a byproduct of DSB formation, to examine the contribution of the DNA damage-responsive kinase Tel1 (ortholog of mammalian ATM) to this regulation in *Saccharomyces cerevisiae*. A *tel1Δ* mutant had globally increased amounts of Spo11-oligonucleotide complexes and altered Spo11-oligonucleotide lengths, consistent with conserved roles for Tel1 in control of DSB number and processing. A kinase-dead *tel1* mutation also increased Spo11-oligonucleotide levels, but mutating known Tel1 phosphotargets on Hop1 and Rec114 did not. Deep sequencing of Spo11 oligonucleotides from *tel1Δ* mutants demonstrated that Tel1 shapes the nonrandom DSB distribution in ways that are distinct but partially overlapping with previously described contributions of the recombination regulator Zip3. Finally, we uncover a context-dependent role for Tel1 in hotspot competition, in which an artificial DSB hotspot inhibits nearby hotspots. Evidence for Tel1-dependent competition involving strong natural hotspots is also provided.

## Introduction

To balance the positive (chromosome segregation promoting) and negative (mutagenic) roles of meiotic DSBs, cells possess overlapping, homeostatic pathways that control the number and repair of DSBs and in doing so shape the recombination distribution (Keeney et al, 2014; Cooper et al, 2016). One such pathway involves the negative regulation of break formation by the DSB-responsive serine/threonine kinase ATM/Tel1 (<u>a</u>taxia telangiectasia <u>m</u>utated in mammals; <u>tel</u>omere maintenance in S. *cerevisiae)*. ATM/Tel1 controls cell-cycle checkpoints and promotes DNA repair in somatic cells and is essential for meiosis in mammals (Shiloh & Ziv, 2013).

Testes from *Atm*^−/−^ mice display a large increase in DSB numbers as indicated by a >10-fold rise in the amount of covalent SpO11-oligonucleotide (oligo) complexes, a quantitative byproduct of meiotic DSB formation (Lange et al, 2011). In *Drosophila melanogaster* oocytes, mutation of ATM elevates numbers of DSB-associated γ-*H2AV* foci (Joyce et al, 2011). And in S. *cerevisiae,* Tel1 loss increases recombination at the *HIS4LEU2* hotspot (Zhang et al, 2011) and genome-wide (Anderson et al, 2015). In a *rad50S* background, in which unrepaired DSBs accumulate, more DSBs are observed at *HIS4LEU2* and on at least one whole chromosome (Carballo et al, 2013; Garcia et al, 2015), although such increases were not apparent on all chromosomes (Argunhan et al, 2013; Blitzblau & Hochwagen, 2013; Garcia et al, 2015). Recently, Tel1 was shown to suppress formation of multiple DSBs nearby on the same DNA molecule (Garcia et al, 2015). In wild-type cells, coincident DSB formation across 70-100 kb domains occurs less frequently than expected from population-average DSB frequencies. However, multiple DSBs cluster on the same chromatid in the absence of Tel1. This mode of ATM/Tel1 action may also ensure that DSBs rarely form at the same location on both sister chromatids (Lange et al, 2011; Anderson et al, 2015; Garcia et al, 2015). Together, these findings revealed that ATM/Tel1 negatively regulates meiotic DSB numbers in many species. The spatial patterning of this regulation is a key but as yet poorly understood feature.

A primary goal of this study was to determine how yeast Tel1 affects the genome-wide number and distribution of DSBs. The DSB “landscape” is defined by complex hierarchical influences operating over different size scales (Pan et al, 2011; Lam & Keeney, 2015a; Cooper et al, 2016). Some genomic regions, encompassing tens to hundreds of kb, experience more or less DSB formation than others, and within such DSB-“hot” and DSB-“cold” domains, DSBs are particularly enriched in small (typically <250 bp) regions known as “hotspots”. In S. *cerevisiae,* most hotspots correspond to evolutionarily conserved nucleosome-depleted promoters (Lam & Keeney, 2015b). However, the factors shaping the DSB landscape at larger size scales remain poorly understood, particularly those factors related to the intersecting feedback mechanisms that regulate Spo11 activity (Keeney et al, 2014; Cooper et al, 2016). To address these issues, we quantified Spo11-oligo complexes and mapped the genome-wide changes in DSB locations upon loss of Tel1. The data provide insight into characteristic genome-wide features of Tel1 loss and relationships among the negative feedback pathways that restrain DSB formation.

## Results

### Increased Spo11-oligo numbers in *tellΔ*

In yeast, global DSB levels are often estimated by Southern blotting of chromosomes separated on pulsed-field gels, often using repair-defective mutant backgrounds. Such studies of *tel1Δ* mutants gave conflicting results (Argunhan et al, 2013; Blitzblau & Hochwagen, 2013; Carballo et al, 2013; Garcia et al, 2015), so we assayed Spo11-oligo levels instead. Spo11 makes a DSB by forming covalent protein-DNA complexes that are then endonucleolytically cleaved to release Spo11 bound to a DNA oligo, with two Spo11-oligo complexes generated from each DSB (**Fig. EV1**) (Neale et al, 2005). Because Spo11-oligo complexes are a direct byproduct of DSB formation, they can be used to quantify relative DSB levels in an otherwise wild-type background, mitigating confounding effects of repair-defective mutations (Lange et al, 2011; Thacker et al, 2014).

Whole-cell extracts were prepared from equivalent numbers of wild-type and *tel1Δ* cells harvested at different times after meiotic induction. Phenotypically normal protein-A-tagged Spo11 was affinity-purified, 3′ ends of Spo11 oligos were radiolabeled using terminal deoxynucleotidyl transferase and [α-^32^P] dCTP, and Spo11-oligo complexes were separated on SDS-PAGE gels and detected by phosphorimager (**Fig. 1A, top**). A western blot for total Spo11 confirms that Spo11 protein expression is induced to a similar extent in wild-type and *tel1Δ* cultures (**Fig. 1A, bottom**). Wild-type cells displayed the typical kinetics of appearance and disappearance of Spo11-oligo complexes (**Fig. 1A, B**), previously shown to coincide with DSB formation (Neale et al, 2005; Thacker et al, 2014). In the *tell A* mutant, Spo11-oligo complexes first appeared with similar timing and levels as wild type but continued to accumulate, reaching a maximum at 5 h that was 2.2-fold higher on average than in wild type (**Fig. 1A, B**). Increased numbers of Spo11-oligo complexes were also apparent in the western blot detecting total Spo11: a slower-migrating band was now apparent that co-migrated with the upper radioactive signal. (The species corresponding to the lower radioactive signal was masked on the western blot by the signal for free Spo11). These results confirm that loss of Tel1 increases DSBs globally.

**Fig. 1.**
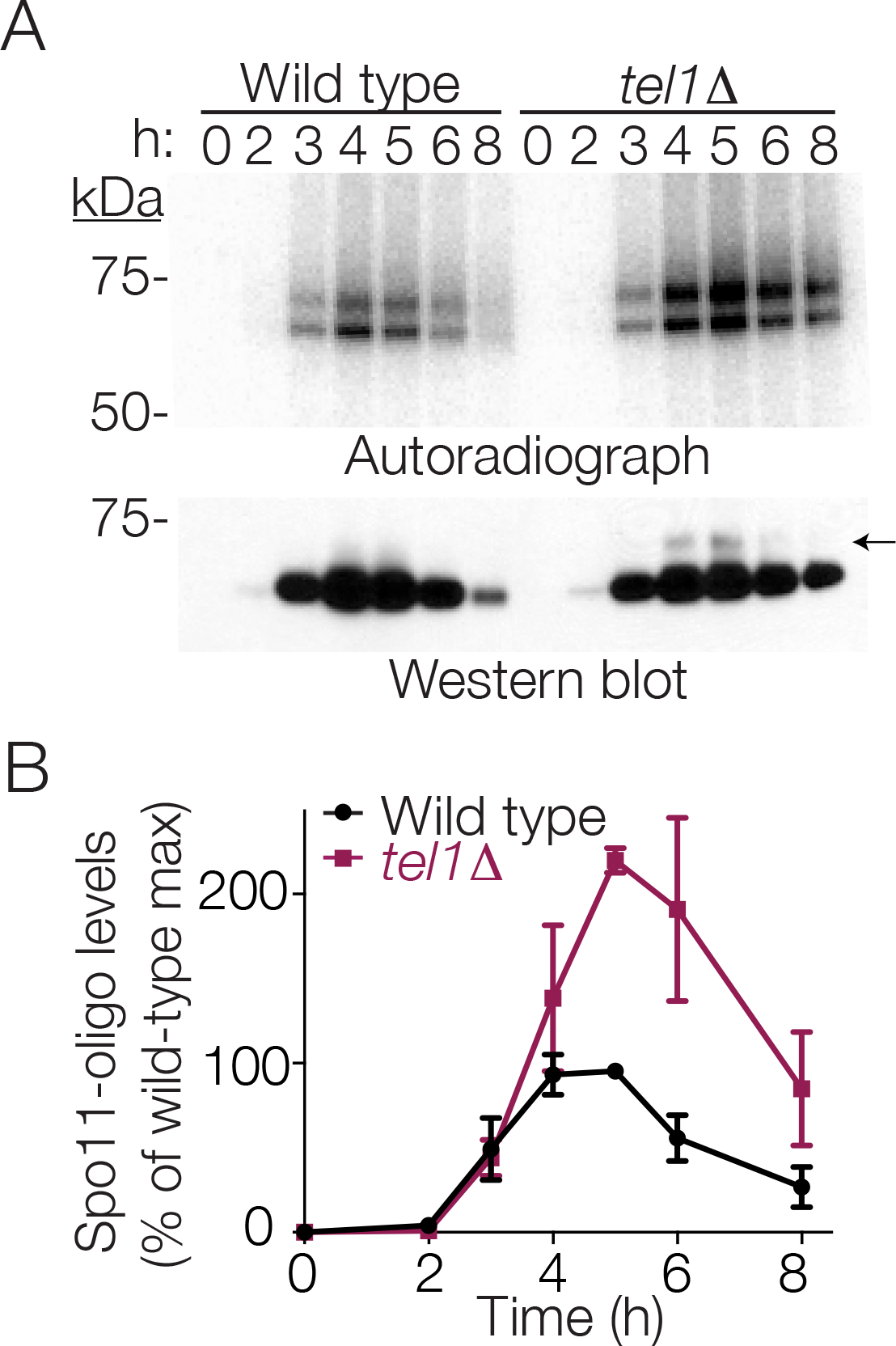
Elevated levels of Spo11-oligo complexes in *tellΔ* mutants. A: Representative time course. Autoradiograph of SDS-PAGE gel (top) shows radiolabeled Spo11-oligo complexes at the indicated times in sporulation medium. Anti-protein A western blot (bottom) monitors total Spo11. Note that most signal in the western blot is free Spo11, i.e., not bound to oligos. The arrow indicates the small amount of Spo11 protein that co-migrates with slower migrating Spo11-oligo species, visible in the *tellΔ* sample because of increased DSB numbers. B: Quantification of radiolabeled Spo11-oligo complexes in *tellΔ* cells, relative to wild-type cultures collected in parallel. Mean ± s.d. for three pairs of cultures is shown.

### Roles of Tell kinase activity and Tell phosphoryation targets

An unresolved question is whether negative regulation of DSB numbers by ATM/Tel1 depends on its kinase activity and, if so, what the relevant phosphorylation targets are. To address these issues, we first asked if a known *tel1* kinase-dead mutation *(“tel1-kd”)* (Ma & Greider, 2009) increased DSBs. Indeed, Spo11-oligo complexes were elevated in *tel1-kd* cultures to an extent similar to that in *tel1Δ* mutants (**Fig. 2A**). These results suggest that Tel1 can suppress DSB formation via the phosphorylation of one or more target proteins.

A candidate target is Rec114, which is required for DSB formation and bears matches to the S/TQ phosphorylation motif for Tel1 and its related kinase Mec1 (Carballo et al, 2013). Mec1 phosphorylates Rec114 in vitro (Tel1 was not tested); Mec1 and Tel1 are redundantly required for DSB-dependent phosphorylation of Rec114 in vivo; and mutation of eight S/TQ sequences in Rec114 to Aq *(“rec1l4-8A”)* yielded indirect evidence of increased and/or earlier DSB formation (Carballo et al, 2013). We therefore asked if *rec114-8A* phenocopies absence of Tel1. Strikingly, although the *rec114-8A* mutant showed a slight increase in Spo11-oligo complexes at late time points, this was substantially less than in the *tel1Δ* or *tel1-kd* mutants (**Fig. 2B**). We also combined *rec114-8A* with mutation of three S/TQ residues of Hop1, another known Tel1 and Mec1 target (known as the *hop1-scd* allele) (Carballo et al, 2008). However, even the small increase seen at late times with *rec114-8A* alone was not apparent in the *rec114-8A hop1-scd* double mutant (**Fig. 2C**). We infer that Rec114 and Hop1 phosphorylation are largely dispensable for Tel1-mediated DSB suppression.

**Fig. 2.**
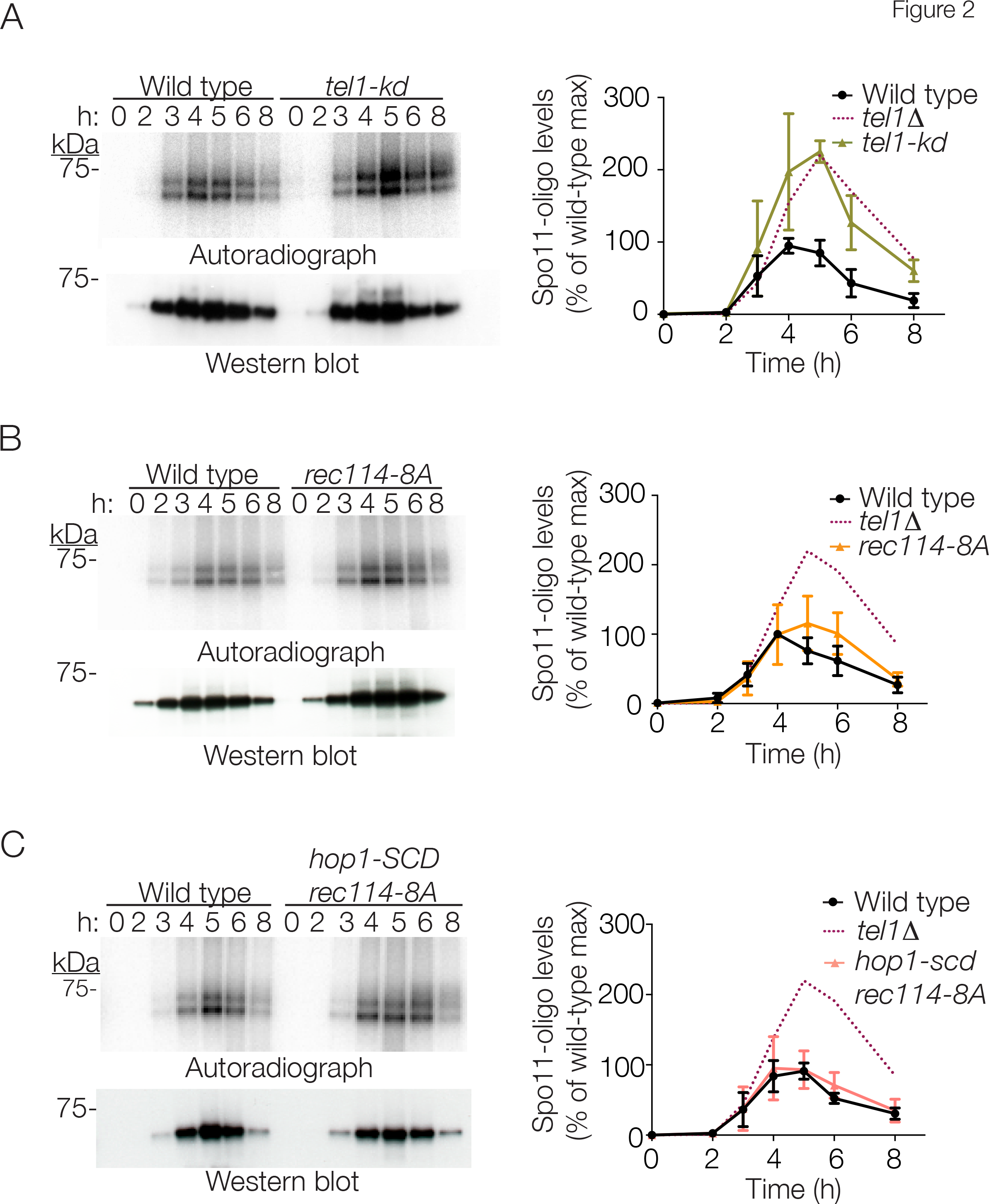
Spo11-oligo complexes in *tel1-kd, rec114-8A* and *hop1-scd mutants*. Representative time courses and quantification of (A) two, (B) four and (C) six independent mutant cultures processed in parallel with (A) three, (B) four and (C) four independent wild-type cultures are shown as in **Fig. 1**. To aid comparison, *tellΔ* data are duplicated from **Fig. 1**.

### Altered Spo11-oligo sizes in *tell* mutants

Most Spo11-oligo complexes are in two size classes that differ in the lengths of the attached oligos (Neale et al, 2005) (**Fig. 1A**). The ratio between these classes appeared altered relative to wild type in the *tel1Δ* mutant, with more of the higher molecular weight species (**Fig. 1A**). To examine this change, Spo11 oligos were separated on a sequencing gel (**Fig. 3A**). Consistent with the SDS-PAGE, *tel1Δ* showed an increase in oligos >25 nt long relative to those <20 nt, including increased abundance of very large oligos near the top of the gel (**Fig. 3A, B**). The distribution of larger oligos was also altered: a modest shoulder at ~35 nt in wild type became more prominent in *tel1Δ*, and even longer oligos (from ~40 to ~70 nt) displayed a more pronounced banding pattern in the mutant (**Fig. 3A, B**).

**Fig. 3.**
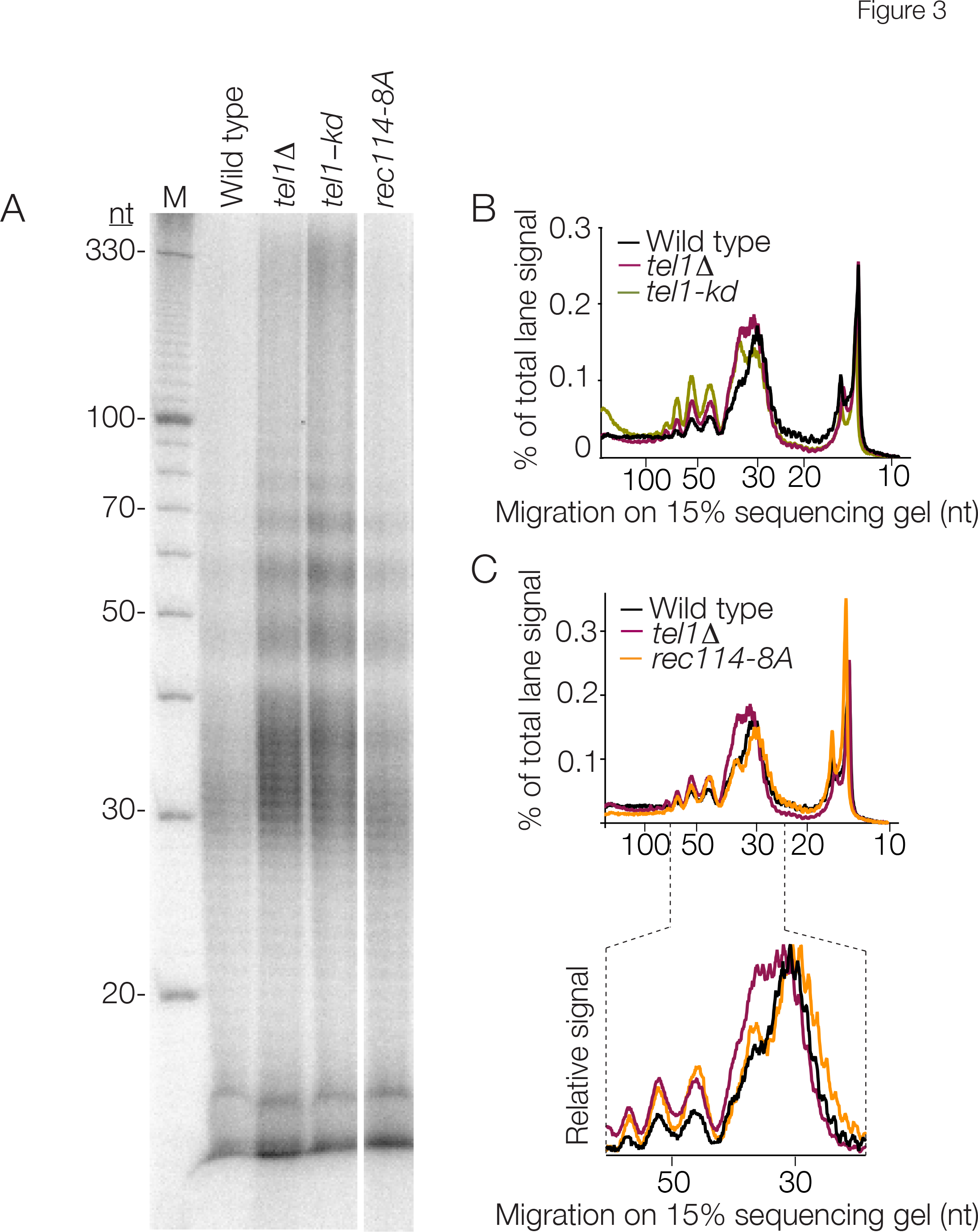
Sizes of Spo11 oligos in Tel1 pathway mutants. A: Radiolabeled, deproteinized Spo11 oligos separated on a 15% denaturing polyacrylamide sequencing gel. Lanes from a single gel are shown; an intervening lane is omitted. M, 10 nt ladder. Note that oligos from lower-molecular-weight Spo11-oligo complexes are underrepresented because most of the shortest oligos (less than ~12 nt) are lost upon ethanol precipitation. B, C: Lane profiles from panel A.

The *tel1-kd* mutant was similar with respect to increased abundance of longer Spo11 oligos (those >25 nt relative to those <20 nt), the more prominent population at ~35 nt, and the banding at ~40-70 nt (**Fig. 3A, B**). Interestingly, these alterations were even more extreme in *tel1-kd* than in *tel1Δ,* indicating that presence of a kinase-dead protein does not precisely phenocopy absence of Tel1.

The distribution of oligo sizes was also altered in the *rec114-8A* mutant, in similar ways but to a lesser degree than in *tel1Δ* (**Fig. 3A,C**). This finding is consistent with Rec114 phosphorylation contributing to regulation of the formation and/or processing of DSBs, but the results also reinforce the conclusion that Rec114 is not the sole Tel1 substrate important for DSB control.

### The DSB landscape responds differently to Tel1 loss at different stages of meiosis

Having confirmed that Tel1 regulates DSB numbers, we investigated how it shapes the DSB distribution. Analysis of recombination products (Anderson et al, 2015) supported an earlier hypothesis that Tel1 shapes the DSB landscape (Lange et al, 2011), but recombination maps are indirect and low-spatial-resolution indicators of DSB positions. We deep-sequenced Spo11 oligos purified from wild-type and *tel1Δ* cultures at 4 h in sporulation conditions, when DSBs are maximal in wild type and the *tel1Δ* mutant has only slightly elevated total Spo11-oligo levels, and at 6 h, when the *tel1Δ* mutant has substantially elevated Spo11 oligos (**Fig. 1B** and **Table S1**). (Maps were also generated from 5-h time points; these consistently showed patterns intermediate between the 4-h and 6-h time points, so for clarity are not discussed further here.) Two *tel1Δ* biological replicates were highly similar (**Fig. 4A, EV2**) and were averaged together. Wild-type samples displayed little systematic variation over this time window, so maps from six samples (two each from 4, 5, and 6 h) were averaged.

At short (sub-kb) size scales, Spo11 oligos from *tel1Δ* cells mapped preferentially to hotspots highly similar in location and width to those from wild type (**Fig. 4A, EV3A,B**). We conclude that the influence of local chromatin structure on hotspot position and shape is largely unaffected by *tel1Δ,* and that the increase in Spo11-oligo levels is not principally from DSBs in regions that were previously inaccessible to Spo11.

**Fig. 4.**
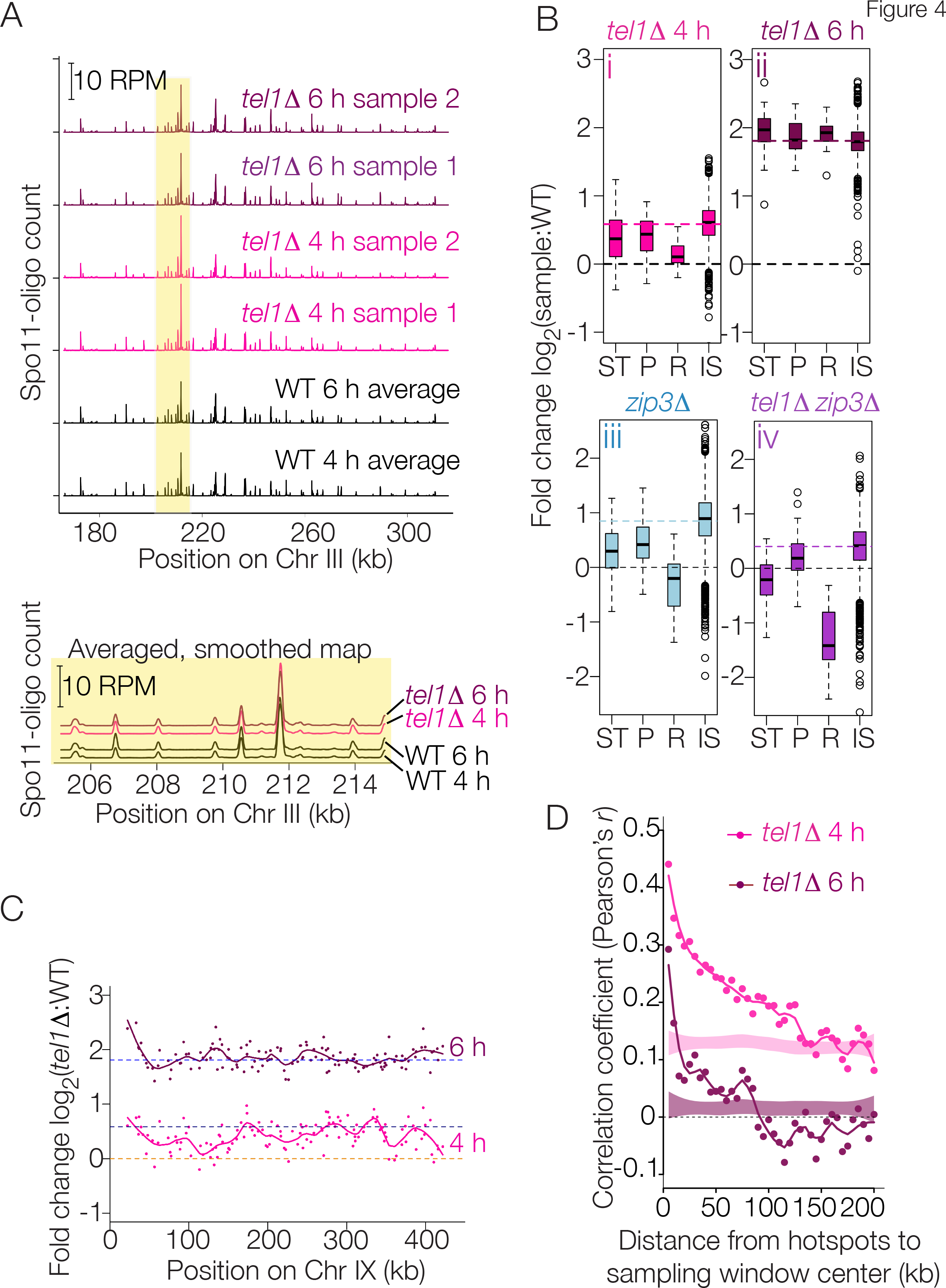
Spo11-oligo landscapes. A: Reproducibility of Spo11-oligo maps. A representative segment from Chr III is shown at top, and a close-up view of averaged biological replicates in the yellow-highlighted region is shown below. Both Spo11-oligo maps were smoothed with a 201-bp Hann window. B: Sub-chromosomal domains. Boxplots show the log-fold change in Spo11-oligo counts in each mutant relative to wild type for hotspots in the indicated genomic regions: ST, sub-telomeric, 20 kb at each chromosome end (62 hotspots); P, pericentric, 20 kb centered at each centromere (82 hotspots); R, rDNA-proximal region (Chr XII: 451,577-467,570; 20 hotspots); IS, interstitial, i.e., all other genomic regions. Thick horizontal lines are medians, box edges are the 25th and 75th percentiles, whiskers indicate lowest and highest values within 1.5 fold of the interquartile range, and open circles are outliers. Colored dashed lines show the genome-average fold change for each mutant (1.5 for *tellΔ* 4 h, 3.5 for *tellΔ* 6 h, 1.8 for *zip3Δ,* and 1.3 for *tellΔ zip3Δ)*. Black dashed lines at zero denote no change. C: Regional variation in the response of hotspots to the absence of Tel1. Chr IX is shown as an example. Each point is the log-fold change in Spo11-oligo count relative to wild type. Solid lines are local regression curves; dashed lines indicate genome-wide average fold change or no change. D: Local domains of correlated behavior. Each point compares the log-fold change in hotspots with the change in their neighbors located within a 5-kb window the indicated distance away. Shaded regions show 95% confidence intervals for hotspots randomized within-chromosome; randomized data at 4 h show higher correlation because of the chromosome size effect (see **Fig. 5A**).

When assessed on larger size scales, however, *tel1Δ* showed substantial changes that evolved over time. At 4 h, Spo11-oligo frequencies had increased less than average in hotspots within 20-kb zones at telomeres and centromeres and had increased even less near the ribosomal DNA (rDNA) (**Fig. 4Bi**). At 6 h, in contrast, hotspots in these domains were similar to or higher than genome average, i.e., were all about equally elevated (**Fig. 4Bii**). Thus, these domains, which are suppressed for DSBs in wild type (Pan et al, 2011), are initially somewhat insulated from the elevation in DSB levels caused by Tel1 deficiency, but over time continue to accumulate additional DSBs to reach a similar extent as elsewhere in the genome.

The remaining interstitial hotspots showed variable responses to the *tel1Δ* mutation at both time points (**Fig. 4Bi, 4Bii, 4C**), but with only a modest correlation between the 4 h and 6 h samples (Fig. EV3C; Pearson’s *r* = 0.34). Peaks and valleys in local regression along chromosomes suggested that groups of hotspots respond to the *tel1Δ* mutation in a similar manner (**Fig. 4C**). To test this conclusion, we measured the correlation in response between hotspots and their neighbors. At 4 h, the change in each hotspot’s Spo11-oligo count was positively correlated with the change in its neighbors, with correlation strength decaying with distance but still above randomized controls up to ~125 kb away (**Fig. 4D**). In contrast, the 6-h time point had a weaker starting correlation for close hotspots (≤5 kb) and this correlation decayed to background levels by 15 kb (**Fig. 4D**). Thus, while hotspots appear to respond to absence of Tel1 in a “domainal” fashion early in meiosis, this organization is largely erased as meiosis progresses. We conclude that loss of Tel1 leads to substantial alterations in the DSB landscape, and these alterations evolve over time as cells continue to make DSBs.

### Relationship between Tel1 and the homolog engagement pathway for DSB control

DSB numbers are also elevated in mutants lacking “ZMM” proteins (Thacker et al, 2014), factors needed to complete recombination and synaptonemal complex formation (Lynn et al, 2007). This and findings from other organisms (Wojtasz et al, 2009; Hayashi et al, 2010; Kauppi et al, 2013) have led to the hypothesis that homologous chromosomes that successfully engage one another stop making DSBs (Kauppi et al, 2013; Thacker et al, 2014). This negative feedback regulation, which contributes to DSB spatial patterning (Thacker et al, 2014), is proposed to be distinct from Tel1-mediated DSB control (Keeney et al, 2014). To test this proposal, we compared how the DSB landscape changes in the absence of Tel1 to how it changes in the absence of the ZMM protein Zip3.

In wild type, smaller chromosomes generate more DSBs per kb than larger ones, revealed as an inverse correlation between Spo11-oligo density and chromosome size (Pan et al, 2011). This anticorrelation is lost in *zip3Δ,* indicating a role for ZMM proteins and thus homolog engagement (Thacker et al, 2014) (**Fig. 5A**). The *tel1Δ* 4-h time point displayed an attenuated version of the *zip3Δ* pattern, i.e. a reduced but not absent anticorrelation (**Fig. 5A**). Similarly, the responses of sub-telomeric, pericentric, and rDNA-proximal regions to absence of Tel1 at 4 h were qualitatively similar to those in the *zip3Δ* mutant but in milder form (**Fig. 4Bi and iii**), and there was a positive correlation between the *tel1Δ* 4-h and *zip3Δ* samples for changes at interstitial hotspots (**Fig. EV3C**, *r* = 0.53). The two datasets also displayed similar correlation strengths and length scales for the local correlation in hotspot behavior (**Fig. 5B**).

**Fig. 5.**
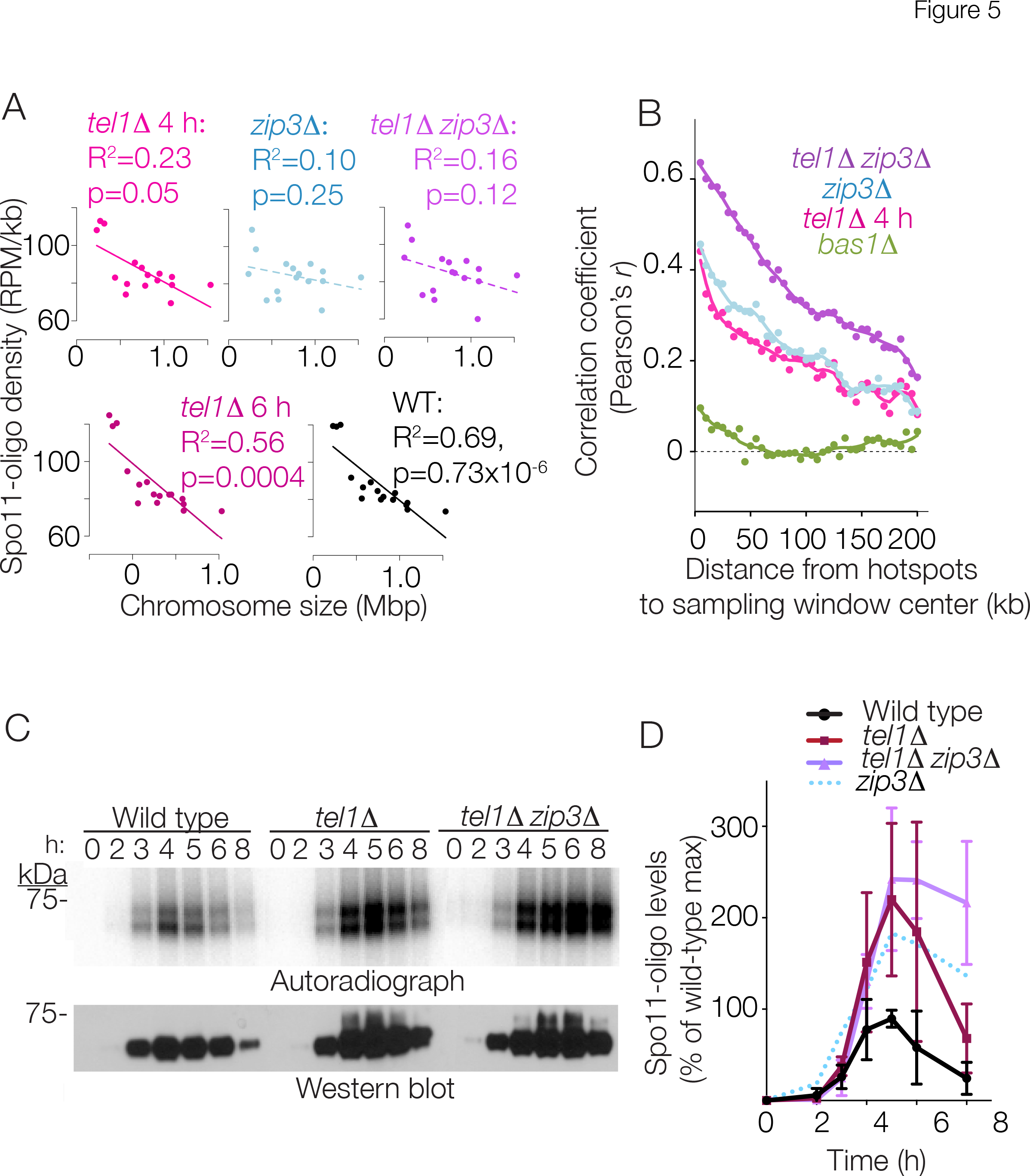
Relationship between Tel1-and Zip3-dependent DSB control pathways. A: Negative correlation between Spo11-oligo density and chromosome size requires Zip3 but not Tel1. *zip3Δ* data are from (Thacker et al, 2014). B: Local domains of correlated behavior (as in **Fig. 4D**). Data for *tellΔ* 4 h are reproduced from **Fig. 4D**. See **Fig. EV4C** for randomized controls. Data from a mutant lacking the transcription factor Bas1 serve as a negative control for this analysis, because this transcription factor affects DSB formation within a subset of individual hotspots rather than coordinately influencing local domains containing multiple hotspots (Zhu & Keeney, 2015). C, D: Spo11-oligo time courses as in **Fig. 1**. Mean ± s.d. for four pairs of cultures is shown. Data from *zip3Δ* are reproduced from (Thacker et al, 2014) for comparison (see also **Fig EV4B**).

However, essentially all similarity with the *zip3Δ* mutant was erased by 6 h in the *tel1Δ* mutant, whether compared in specific sub-chromosomal domains (**Fig. 4B**), in local domains of correlated behavior (**Fig. 4D, 5B**), or across all interstitial hotspots (**Fig. EV3C**, *r* =-0.01). Furthermore, at 6 h the negative correlation between Spo11-oligo density and chromosome size mirrored the pattern in wild type **Fig. 5A**. Thus, Tel1 is dispensable for chromosome size-related control of DSB frequency, although it is needed for timely establishment.

To further characterize the apparent similarities between the *tel1Δ* 4-h and *zip3Δ* DSB landscapes, we examined a *tel1Δ zip3Δ* double mutant. At time points up to ~5 h, the double mutant yielded levels of Spo11-oligo complexes that were indistinguishable from the *tel1*Δ single mutant, i.e., with maximum elevated ~2.4-fold relative to wild type (**Fig. 5C,D**). For comparison, we previously found that the peak was elevated by 1.8-fold in the *zip3Δ* mutant (Thacker et al, 2014) (**Fig. 5D, EV4B**). These findings indicate that loss of both Tel1 and ZMM function does not cause an additive increase in DSB numbers early in meiosis, suggesting at least some degree of overlap between these mechanisms of DSB control. At later time points, the *tel1Δ zip3Δ* double mutant maintained Spo11-oligo complexes at high levels (**Fig. 5C,D**). This is unlike the *tel1Δ* single mutant in which Spo11-oligo complexes decreased over time, but is more reminiscent of the pattern in the *zip3Δ* single mutant (**Fig. EV4B**).

Analysis of Spo11-oligo maps from the *tel1Δ zip3Δ* double mutant at 4 h in meiosis (**Fig. EV4A**) indicated that the epistasis relationships between the *tel1Δ* and *zip3Δ* mutations are complex. There was no longer a residual anti-correlation between Spo11-oligo density and chromosome size in the double mutant, indistinguishable from the *zip3Δ* single mutant (**Fig. 5A**). Similarly, the double mutant more closely mirrored the stronger patterns seen in the *zip3Δ* single mutant for hotspots in the sub-telomeric and rDNA-proximal regions, but less so in pericentric regions (**Fig. 4Biv**). Thus, *zip3Δ* is epistatic to *tel1Δ* for these features of the DSB landscape.

For other features, however, Spo11-oligo maps revealed largely non-epistatic relations instead. For interstitial hotspots, the log-fold change of Spo11-oligo counts in the double mutant correlated best with the *zip3Δ* single mutant (**Fig. EV3C**, *r* = 0.79), so most of the change in the double mutant can be accounted for by the *zip3Δ* mutation. But there was also a stronger correlation of the *tel1Δ* 4-h dataset with *tel1Δ zip3Δ* than with the *zip3Δ* single mutant (**Fig. EV3C**, *r* = 0.62 vs. 0.53), and the *tel1Δ* 6-h dataset now displayed a modest correlation with the *tel1A zip3A* map that was not seen with the *zip3Δ* map (**Fig. EV3C**, *r* = 0.18 vs.-0.01). These observations suggest that the DSB landscape in the double mutant combines the changes shared in common between the two single mutants with features that are unique to either *tel1Δ* or *zip3Δ*. Furthermore, neighboring hotspots displayed more strongly correlated behavior that spread over longer distances (up to ~200 kb) in the *tel1Δ zip3Δ* double mutant than in either single mutant (**Fig. 5B**). This suggests that effects of *tel1Δ* or *zip3Δ* mutations are additive (non-epistatic) for this feature of the DSB landscape. Possible explanations for this complex behavior are provided in the Discussion.

### Context-dependent contribution of Tel1 to hotspot competition

Inserting a strong artificial hotspot suppresses DSB formation nearby (Wu & Lichten, 1995; Xu & Kleckner, 1995), a phenomenon called hotspot competition (Keeney et al, 2014). This suppressive effect decays with distance but has been documented to extend ≥60 kb (Wu & Lichten, 1995; Robine et al, 2007). Site-specific endonucleolytic DSBs also suppress Spo11-generated DSBs nearby (Fukuda et al, 2008); if this works by the same mechanism, it implies that hotspot competition involves an inhibitory signal that spreads after DSB formation. Tel1 is a plausible candidate for the generator of this signal given its role in DSB interference (Garcia et al, 2015). To test for Tel1 involvement, we inserted the *arg4-URA3* artificial hotspot (Wu & Lichten, 1995) at either of two positions in the genomes of otherwise wild-type or *tel1Δ* cells: at *HIS4* on Chr III or *POR2* on Chr IX. We then examined the effects of the hotspot insertions in cis by compiling Spo11-oligo maps from 4-h meiotic cultures.

As previously demonstrated (Wu & Lichten, 1995), hotspot insertion on Chr III in a *TEL1+* background caused a substantial drop in DSB formation nearby: Spo11-oligo density was decreased by four-fold or more in hotspots flanking the insertion site (**Fig. 6A, 6B**). Suppression decayed with distance, extending ~60 kb to the left (nearly to the telomere) and ~130 kb to the right, extending past the centromere, and even spanning, albeit more weakly, the DSB-cold domain from *CEN3* to *MAT*. In *tel1Δ,* DSBs were suppressed to a similar degree from the insertion site to the left and from the insertion site to the centromere, but the *CEN3-MAT* region was no longer suppressed (**Fig. 6A, 6B**). We conclude that for this insertion site, Tel1 is dispensable for hotspot competition over distances up to ~60 kb, but may contribute to a form of hotspot competition that operates over longer range.

**Fig. 6.**
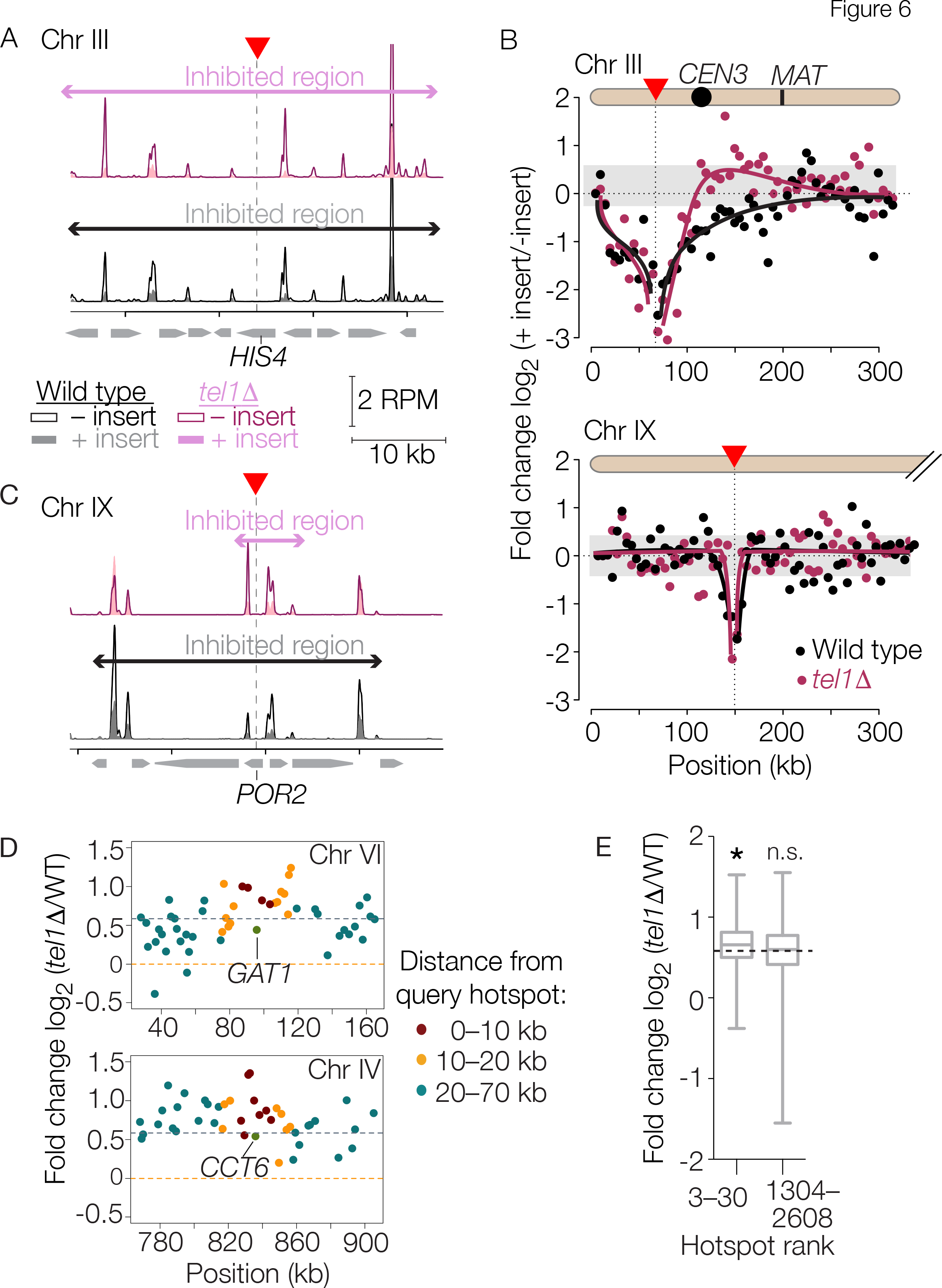
Hotspot competition. A-C: Effects of artificial hotspot insertions. A hotspot construct containing *arg4* and *URA3* on a bacterial plasmid backbone (Wu & Lichten, 1995) was inserted within the *HIS4* gene on Chr III or the *POR2* gene on Chr IX (red triangles and vertical dotted lines). (A,C) Spo11 oligo maps (smoothed with a 201-bp Hann sliding window) are shown for wild type and *tel1Δ* strains with and without the insertion. (B) Each point shows, for each 5-kb segment moving outward from the insertion points, the log-fold change in Spo11-oligo count caused by presence of the artificial hotspot insertion. Curves were drawn manually to highlight trends in the data. Gray shading encloses ± 1 standard deviation for all 5-kb bins genome wide (excepting Chr III and IX), to illustrate range of sampling noise in this analysis. D,E: Signature consistent with Tel1-dependent hotspot competition around the strongest natural hotspots. In panel D, each point displays the log-fold change in Spo11-oligo count (4-h *tel1Δ* vs. wild type) for hotspots located the indicated distance away from either the *GAT1* (top) or *CCT6* (bottom) promoter hotspots. Horizontal dashed lines indicate genome-average change or no change. In panel E, boxplots summarize the log-fold change for hotspots located within 10 kb of the next 28 hottest hotspots (rank numbers 3-30) or within 10 kb of the middle third of hotspots (rank numbers 1304-2608). Asterisk, p = 0.0011 for a one-sided Wilcoxon rank sum comparison to all hotspots genome-wide; n.s., not significant (p = 0.61). Horizontal dashed line indicates genome-average change. Boxplots are as defined in **Fig. 4B**.

Chr IX gave a strikingly different result. In the *TEL1+* background, this hotspot insertion caused a substantial drop in Spo11-oligo counts for hotspots immediately flanking the insertion site, yielding a similar magnitude of suppression as on Chr III (**Fig. 6B, 6C**). However, the suppressive effect covered a much smaller zone, only ~20 kb to left and ~15 kb to right. The more limited distance of suppression may be in part a consequence of the artificial hotspot being weaker at the Chr IX location (**Fig. EV5**). Context dependence for DSB frequency within this insert was documented previously (Borde et al, 1999). In the *tel1Δ* background, the hotspots closest to the insertion (<2 kb away) were still suppressed, but the next hotspots further away (≥5 kb) were unaffected by the insertion (**Fig. 6b, 6C**). Thus, the zone of hotspot competition is much smaller at this location and Tel1 is essential for DSB suppression beyond the most immediate vicinity of the new hotspot. More broadly, these results show that hotspot competition is context dependent, in terms both of the distance over which it works and its dependence on Tel1.

A prior study found that a natural hotspot in the *HIS4* promoter can exert a suppressive effect on an artificial one inserted nearby (Fan et al, 1997), but another study found no evidence of competition between two natural hotspots near *HIS2* (Bullard et al, 1996). It has thus been unclear whether hotspot competition is unique to artificial hotspots. We reasoned that, if this phenomenon does indeed apply to natural hotspots and if it is at least partially Tel1-dependent like the insertion on Chr IX, then absence of Tel1 should cause greater than average increase in DSBs for hotspots located near strong hotspots, because weak hotspots should be more highly suppressed by Tel1 signals emanating from their stronger neighbors.

To test this prediction, we examined *GAT1,* the strongest natural hotspot in the SK1 strain background (Pan et al, 2011). At 4 h in the *tel1Δ* mutant, this hotspot displayed a Spo11-oligo count that was slightly below the genome-average increase (**Fig. 6D, Ev6A**). In contrast, nearly all of its neighbors within ~10-20 kb on either side experienced above-average increases in Spo11-oligo levels; hotspots further away were not consistently increased above genome average (**Fig. 6D**). Similarly, most hotspots in the vicinity of another exceptionally strong natural hotspot, *CCT6*, also increased more than average (**Fig. 6D, EV6B**), as did hotspots located up to ~10 kb away from the next 28 hottest hotspots considered as a group (**Fig. 6E**). This pattern meets the prediction for Tel1-dependent competition between strong hotspots and their weaker neighbors. Notably, however, we detected no reliable hotspot-competition signature for hotspots that are less active than this elite group (**Fig. 6E**).

Taken together, these findings support the hypothesis that hotspot competition is a general phenomenon that is substantially but not fully Tel1 dependent, that usually extends for only short distances, that is readily detected only at relatively strong hotspots, and that is quantitatively modest on a population average. It appears that effects of the artificial hotspot closely mimic competition between natural hotspots in terms of distance and Tel1 dependence when inserted on Chr IX, but insertion on Chr III is atypical.

## Discussion

Quantification of Spo11-oligo complexes reinforces the conclusion that more DSBs form when Tel1 is missing in S. *cerevisiae,* which was indicated by many studies (Zhang et al, 2011; Carballo et al, 2013; Anderson et al, 2015; Garcia et al, 2015) but not all (Argunhan et al, 2013; Blitzblau & Hochwagen, 2013). However, although it is now clear that Tel1/ATM has an evolutionarily conserved function in regulating DSB numbers, the mechanism remains poorly understood. Tel1 kinase activity appears to be critical, but the relevant phosphotarget(s) is unknown. Prior studies implicated Rec114 phosphorylation (Carballo et al, 2013), but we found that mutating the presumed S/TQ phosphoacceptor sites mimicked absence of Tel1 only weakly if at all. It is therefore likely that dSb suppression proceeds via another phosphotarget(s) and does so either exclusively of or redundantly with the tested S/TQ motifs of Rec114 and Hop1. We favor the redundancy interpretation because it provides a straightforward explanation for the modest increase in DSB levels and partial alteration of Spo11-oligo sizes in the *rec114-8A* mutant (which together indicate that Rec114 is indeed a functional target of Tel1) and the fact that phosphomimetic mutations at Rec114 S/TQ sites substantially reduce DSBs (Carballo et al, 2013) (which may indicate that Rec114 phosphorylation is sufficient but not necessary for DSB inhibition). Identifying the other phosphotarget(s) will be an important next step.

We find that Tel1 also has a conserved role in modulating the lengths of Spo11 oligos. *Atm*^−/−^ spermatocytes have alterations in SPO11 oligo sizes that are strikingly similar to those observed in *tel1Δ* and *tel1-kd,* namely, fewer short oligos relative to long ones, a more pronounced banding pattern for oligos in the ~40-70 nt range, and an increase in abundance of very long oligos (Lange et al, 2011). One model to explain these findings is that Tel1 acts early in meiotic DSB processing, and indeed Tel1 phosphorylates proteins necessary for Spo11 removal from DNA ends, such as Mre11 and Sae2 (Cartagena-Lirola et al, 2006; Terasawa et al, 2008). Thus, altered oligo sizes may reflect changes in the positions of Mre11-dependent endonucleolytic cleavage (Cannavo & Cejka, 2014) and/or decreased 3′→5′ exonuclease activity of Mre11 (Garcia et al, 2011). However, an alternative, non-exclusive model is that altered Spo11-oligo sizes arise from adjacent DSBs on the same DNA molecule (Garcia et al, 2015). In this case, the size changes are a consequence of aberrant control of dSb numbers rather than (or in addition to) a change in nucleolytic DSB processing. This model more readily accounts for the apparent contribution of Rec114 phosphorylation, and for the observation that reducing DSB numbers (by *Spo11*^+/−^ heterozygosity) in an *Atm*^−/−^ background partially suppresses the changes in SPO11-oligo sizes (Lange et al, 2011). In either model, it is unclear why presence of catalytically inactive Tel1 should exacerbate the changes. In mice, an ATM kinase-dead mutation causes embryonic lethality whereas deletion mutants are viable, providing precedent for critical functional differences between the two types of allele (Daniel et al, 2012; Yamamoto et al, 2012).

Because Tel1/ATM is activated in the vicinity of DSBs, it was argued that it would contribute to spatial patterning of DSBs across the genome (Lange et al, 2011). Spo11-oligo mapping affirmed this prediction, but also unexpectedly demonstrated that the effect of Tel1 absence differs over the course of meiosis. At 4 h, the DSB landscape in *tel1Δ* resembled that in *zip3Δ,* but by 6 h that similarity was largely gone and the *tel1Δ* mutant instead displayed a unique but modest set of differences from wild type. We consider two straightforward ways to rationalize the similarity between the early *tel1Δ* map and the *zip3Δ* map. First, subchromosomal domains that are more or less sensitive to Tel1-mediated DSB suppression might also be correspondingly more or less sensitive to ZMM-dependent suppression. For example, domains enriched for Tel1 substrates might be more sensitive to Tel1 regulation; of note, enrichment for Hop1, Red1, and Rec114 is correlated with stronger ZMM-dependent DSB suppression (Thacker et al, 2014). A second, non-exclusive possibility follows from noting that Tel1 promotes inter-homolog bias early in meiosis (Joshi et al, 2015). Meiotic recombination favors the use of the homolog rather than the sister chromatid as the repair partner for DSBs (Hunter & Kleckner, 2001), and this inter-homolog bias depends on Tel1 or its paralog Mec1 (related to mammalian aTr) (Carballo et al, 2008). Tel1 is principally responsible for this function early in meiosis, when DSB levels are relatively low, but it is rendered redundant with Mec1 later, when DSB levels are high (Joshi et al, 2015). We therefore reason that *tel1Δ* mutants, because of reduced inter-homolog bias, experience a delay in homolog engagement and may as a consequence initially display many of the alterations in the DSB landscape that are diagnostic of the ZMM mutation *zip3Δ*. In this scenario, substitution of Mec1 for Tel1 activity eventually overcomes the transient homolog engagement defect. One implication is that the homeostatic compensation that occurs dynamically over the course of meiosis is generally invisible to methods that rely on examining only the final outcome of recombination, e.g., tetrad analysis.

Both models also provide an explanation for the complex epistasis relationship between the *tel1Δ* and *zip3Δ* mutations, in which some measurements such as amount of Spo11-oligo complexes indicate a non-additive (i.e., at least partially epistatic) relationship, whereas other measures, including genome-wide DSB distributions, show additive (non-epistatic) behaviors. Taken together, the results are consistent with the hypothesis that Tel1 and ZMM proteins regulate DSB formation via distinct mechanisms, but also indicate that these pathways for DSB control overlap with one another because of the chromosome-associated proteins that implement the control and/or because both pathways are needed to support proper engagement between homologous chromosomes.

What then are the unique contributions of Tel1 to shaping the DSB landscape? Perhaps the most prominent effect is via Tel1 control of DSB interference, which discourages cutting of the same chromatid more than once within a small distance (Garcia et al, 2015). As pointed out by Neale and colleagues, however, DSB interference manifests itself principally at the level of individual cells, and is thus predicted to leave relatively little footprint in the population-average DSB map (Garcia et al, 2015). Consistent with this view, the 6-h *tel1Δ* Spo11-oligo map displayed only relatively modest changes, indicating that the net effect of Tel1 absence is that the extra DSBs end up accumulating largely in the same places as they do in wild type. Redundancy with Mec1, especially later in meiosis (Joshi et al, 2015) may also contribute to Tel1 having a relatively modest net effect on the DSB landscape. Interestingly, the major Tel1 effects at the population level appeared to be highly local (within several kb), as judged by the short distance over which correlated changes were observed in the *tel1Δ* mutant at 6 h. It is possible that this pattern reflects a major function for Tel1 in controlling DSBs within single chromatin loops and, to a lesser extent, between adjacent loops, as previously suggested (Garcia et al, 2015).

Our findings also reveal a function for Tel1 in the phenomenon of hotspot competition. The artificial hotspot insertion on Chr IX showed short-range competition with adjacent hotspots that was independent of Tel1 for hotspots within a few kb, but highly dependent on Tel1 for distances beyond that. Neighbors of the very hottest natural hotspots showed changes in the *tel1Δ* mutant that were consistent with similar short-range Tel1-dependent competition, suggesting this phenomenon is not restricted to artificial hotspots. Weaker natural hotspots failed to show evidence of such Tel1-dependent competition, however. This may indicate that Tel1 dependence varies between genomic locations (also supported by results of hotspot insertion on Chr III), and/or that a hotspot with relatively high DSB frequency is necessary to uncover this behavior (Zhang et al, 2011; Zhu & Keeney, 2015; Cooper et al, 2016). This in turn may explain a prior study in which medium-strength natural hotspots failed to display competition (Bullard et al, 1996).

Some of the earliest studies of hotspot competition were on Chr III (Wu & Lichten, 1995; Xu & Kleckner, 1995; Fan et al, 1997). Our studies indicate that hotspot competition on this chromosome has unusual features with respect to Tel1-dependence and the distance over which it spreads. In contrast to hotspot competition, Tel1-dependent DSB interference is readily detectable on Chr III (Garcia et al, 2015); this difference in Tel1 dependence suggests that hotspot competition on this chromosome is not simply a consequence of DSB interference. DSB formation on Chr III is unusual in other regards as well, including a large DSB-suppressed zone from the centromere to the mating-type locus and lack of dependence on the cohesin subunit Rec8, upon which DSB formation is strongly dependent in most other parts of the genome (Klein et al, 1999; Kugou et al, 2009). It is tempting to speculate that this yeast “sex chromosome” has evolved several exceptional behaviors that support its accurate segregation despite a large recombination-suppressed zone that maintains linkage of *MAT* to the centromere.

It has been argued that the elements that shape DSB distrbutions can be divided into “intrinsic” factors (the DSB machinery and its chromatin substrate) and more “extrinsic” (or “reactive”) factors that provide homeostatic feedback regulation (Keeney et al, 2014; Cooper et al, 2016). Our results place Tel1 firmly on the list of such extrinsic factors, and illustrate the complex interplay between Tel1-dependent and ZMM-dependent pathways for controling both the number and distribution of DSBs.

## Materials and Methods

Additional details are provided in Appendix Supplementary Methods.

### Yeast strains

Yeast strains are of the SK1 background (**Table S2**). Tagged Spo11 constructs were as previously described (Thacker et al, 2014). The *hop1-scd* allele *hop1-S298A, S311A, T318A* (Carballo et al, 2008) was provided by R. Cha. The *tel1-kd* allele of Ma and Greider (Ma & Greider, 2009) *(tel1-D2612A,N2617A,D2631A)* was generated via QuickChange mutagenesis on a plasmid-borne fragment of *TEL1* and integrated to replace the endogenous *TEL1* sequence. We recreated the *rec114-8A* allele described by (Carballo et al, 2013) *(S148A, T175A, T179A, S187A, S229A, T238A, S256A, S307A)* using synthetic dNa fragments, and integrated it to replace the endogenous *REC114* gene. The artificial hotspot insertions were created as follows: fragments of the yeast genome containing coding regions from *HIS4* (+726 to +2062 with respect to ATG) or *POR2* (−380 to +1001 with respect to ATG) were cloned into the *EcoRI* sites of pMJ113 (Wu & Lichten, 1995), cut with either *XbaI (HIS4* hotspot) or *BstXI (POR2* hotspot) and transformed into the appropriate SK1 strains. All integrations were confirmed by Southern blotting and/or sequencing of DNA amplified from genomic DNA.

### Spo11-oligo complexes

Cells were cultured and harvested from meiotic time courses as previously described (Thacker et al, 2014). Denaturing extracts were prepared from yeast cells using 10% trichloroacetic acid and agitation in a bead beater, then Spo11-ProtA was affinity-purified on rabbit IgG-coupled magnetic beads. Spo11-oligo complexes were end labeled with [α-^32^P] dCTP and terminal deoxynucleotidyl transferase (TdT), resolved on SDS-PAGE gels, then transferred onto PVDF membranes and visualized by phosphorimager. Western blotting to monitor total Spo11 was done using an anti-Protein A antibody (Sigma). To evaluate lengths of Spo11 oligos, Spo11-oligo complexes were affinity purified and radiolabeled, and then digested with proteinase K. The deproteinized Spo11 oligos were ethanol precipitated, resuspended in formamide dye, and run on a 15% denaturing polyacrylamide sequencing gel with radiolabeled 10-nt DNA ladder as a marker (Life Technologies).

### Spo11-oligo sequencing

Culture conditions and volumes were identical to those described previously (Thacker et al, 2014) except that 200 ml of SPM culture was removed for each time point. Affinity purification of Spo11-ProtA with Fast Flow IgG Sepharose beads (GE) and sequencing library prepration were as previously described (Thacker et al, 2014). Briefly, ~200 fmol of Spo11 oligos were tailed with GTP at their 3′ ends using terminal deoxynucleotidyl transferase, then ligated to a duplex DNA adapter with a 3′ CCCC overhang using T4 RNA ligase 2 (gift of S. Shuman, MSKCC). Complementary strands of Spo11 oligos were synthesized with Klenow fragment of DNA polymerase I, gel-purified on denaturing polyacrylamide gels, then tailed with GTP and ligated to a second duplex adapter. The resulting ligation products were amplified by low cycle numbers of PCR to add Illumina sequencing adapters, and PCR products were gel-purified and sequenced using Illumina HiSeq in the MSKCC Integrated Genomics Operation.

R version 3.1.1 (http://www.r-project.org/) and GraphPad Prism 6.0e were used for analyses. The Illumina reads were mapped to the yeast genome using a pipeline that was described previously (Pan et al, 2011; Thacker et al, 2014) (**Table S1**). Briefly, adaptor sequences were removed from the 5′ and 3′ ends of the reads and the remaining sequence was mapped to the S. *cerevisiae* genome (SGD version June 2008, that is, *sacCer2)* using gmapper-ls (2_1_1b) from the SHRiMP mapping package. Reads that mapped uniquely to the genome (i.e. the vast majority of all reads) were used in subsequent analyses. Raw and processed sequence reads have been deposited in the Gene Expression Omnibus (GEO) repository (http://www.ncbi.nlm.nih.gov/geo/ (accession number pending). Each map was normalized to the total number of reads mapping uniquely to a chromosome (discarding reads mapping to the 2-<plasmid and mitochondrial DNA) and biological replicate maps were averaged.

A new list of hotspots was compiled to take advantage of current availability of a large number of deeply sequenced wild-type Spo11-oligo preparations [this study; X. Zhu, M. van Overbeek and S. Keeney, unpublished; and refs: (Thacker et al, 2014; Zhu & Keeney, 2015)]. These maps were generated using using untagged, HA-tagged, FLAG-tagged, or Protein A-tagged alleles of Spo11, the latter two having been shown to be phenotypically indistinguishable from untagged with respect to DSB levels (Thacker et al, 2014). Hotspots were called on an averaged wild-type Spo11-oligo map using a previously described algorithm (Pan et al, 2011). Briefly, candidate hotspots were first identified as regions where the Spo11-oligo map smoothed with a 201-bp Hann window was >0.2 RPM per bp (i.e., 2.41 times the genome average Spo11-oligo density). Hotspot boundaries within these candidate locations were defined as the leftmost and rightmost coordinates to which at least one Spo11 oligo had mapped. Hotspots separated by <200 bp were merged. In this manner, we compiled 3908 hotspots. This serves as a “definitive” list for wild-type SK1 under these conditions (**Table S3**).

In analyses presented in **Fig. 4B, 4C, 4D, 5B, 6D** and **EV4C** where we evaluated the log-fold change in the heat of hotspots, we assumed a global increase in Spo11-oligo number of 1.5-fold for the *tel1Δ* 4 h time point, 3.5-fold for the *tel1Δ* 6 h time point and 1.3-fold for *tel1Δ zip3Δ,* based on the difference in Spo11-oligo levels relative to wild-type at the indicated time points (**Fig. 1B, 5D**). For ratios, we added a small constant (+20) to numerator and denominator to prevent dividing by zero and to minimize sampling variability for ratios involving small denominators. To examine the spatial correlations between change in hotspot heats as shown in **Fig. 4D, 5B, EV4C** we calculated the Pearson’s correlation coefficient between the log-fold change at each hotspot and the log-fold change for neighboring hotspots within a 5-kb window located from 5 to 200 kb away in steps of 5 kb. Randomized controls for this analysis were generated by randomly reassigning the-log fold change among hotspots within the same chromosome.

## Acknowledgments

We thank A. Viale (MSKCC Integrated Genomics Operation) for sequencing and N. Socci (MSKCC Bioinformatics Core) for mapping the sequencing reads. We thank S. Shuman (MSKCC) for gifts of T4 RNA ligase 2 and M. Lichten (NCI) and R. Cha (Bangor Univ.) for strains and plasmids. This work was supported by National Institutes of Health grants R01 GM058673 and R35 GM118092.

## Author Contributions

NM performed the experiments. NM and SK designed the project, analyzed and interpreted the data, and wrote the manuscript.

## Extended View Figure Legends

**Fig. EV1.**
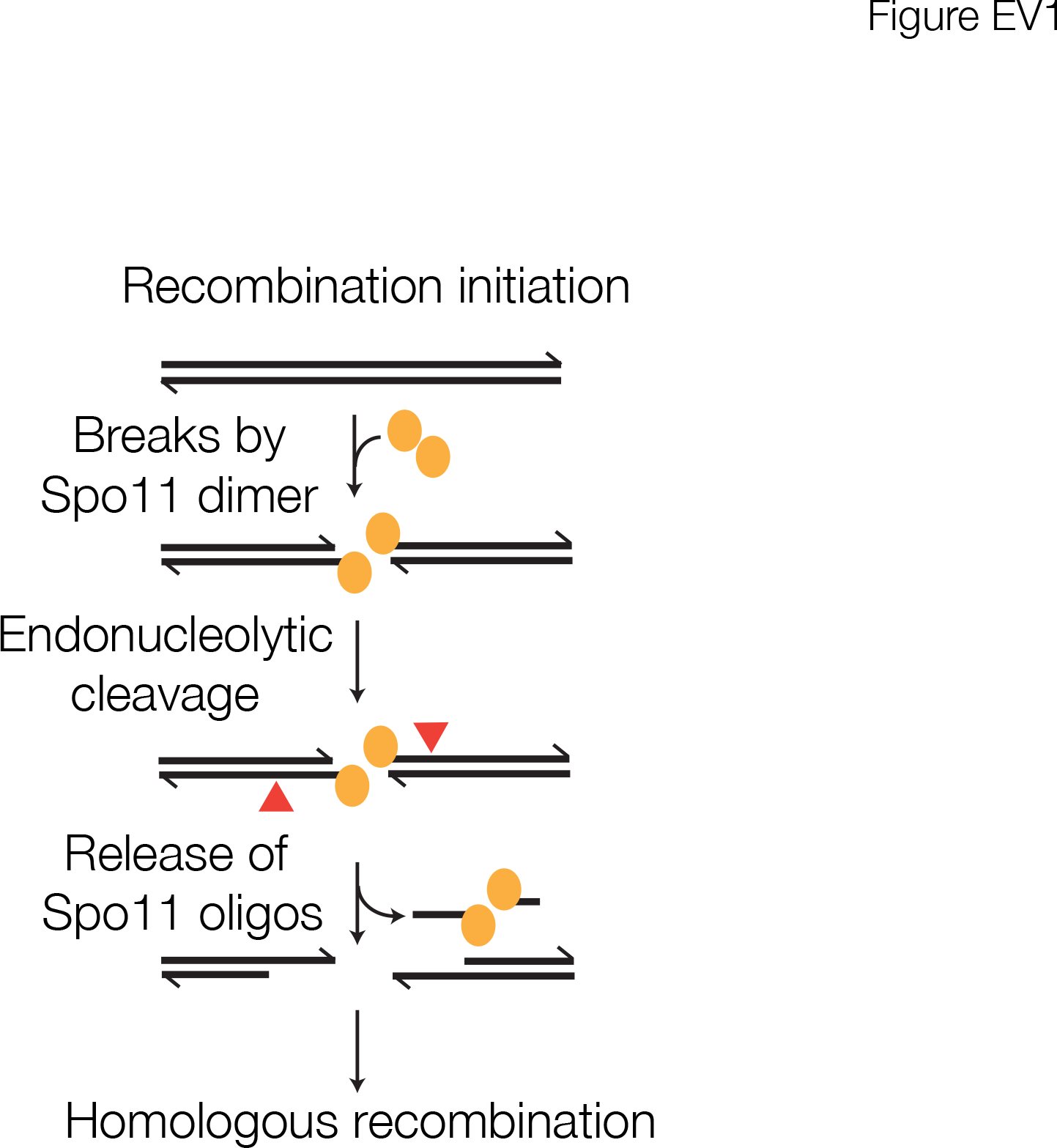
Schematic of recombination initiation pathway. Parallel black lines represent the two strands of double-stranded DNA with arrowheads indicating the 3’ ends. A dimer of Spo11 (yellow ovals) cleaves both strands of the DNA molecule, remaining attached to the 5’ ends of the DSB. Spo11 is released following endonucleolytic cleavage (red triangles) of the strands to which Spo11 is attached.

**Fig. EV2.**
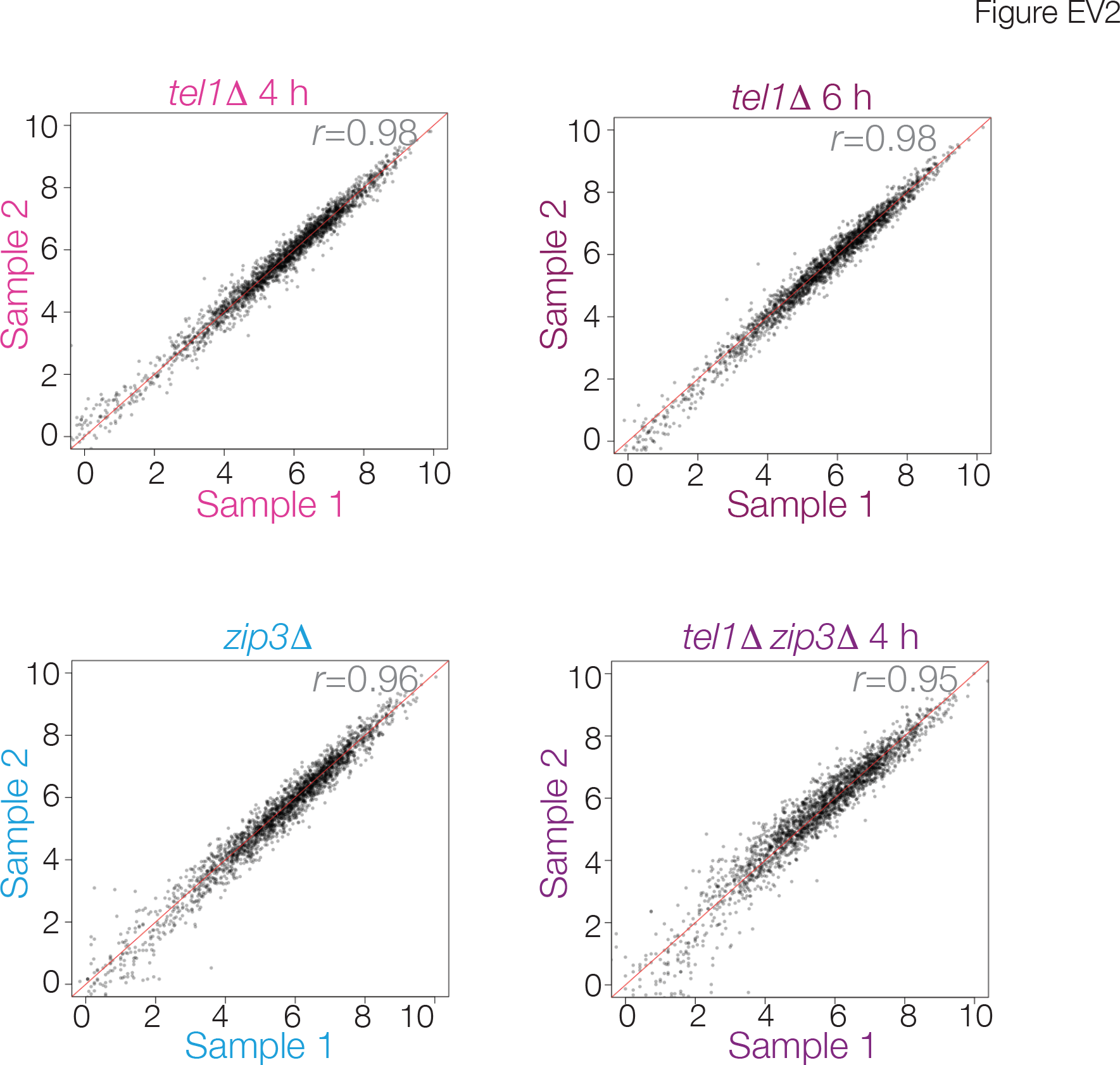
Reproducibility of Spo11-oligo maps. Each point shows the sum of Spo11 oligos (log2) in 5-kb non-overlapping bins across the genome. Biological replicate samples are compared for each genotype and time point indicated. For subsequent analyses, the replicate samples were averaged. Pearson’s correlation values are shown on the top right. Diagonal red line has a slope of 1. The *zip3Δ* samples are from (Thacker et al, 2014).

**Fig. EV3.**
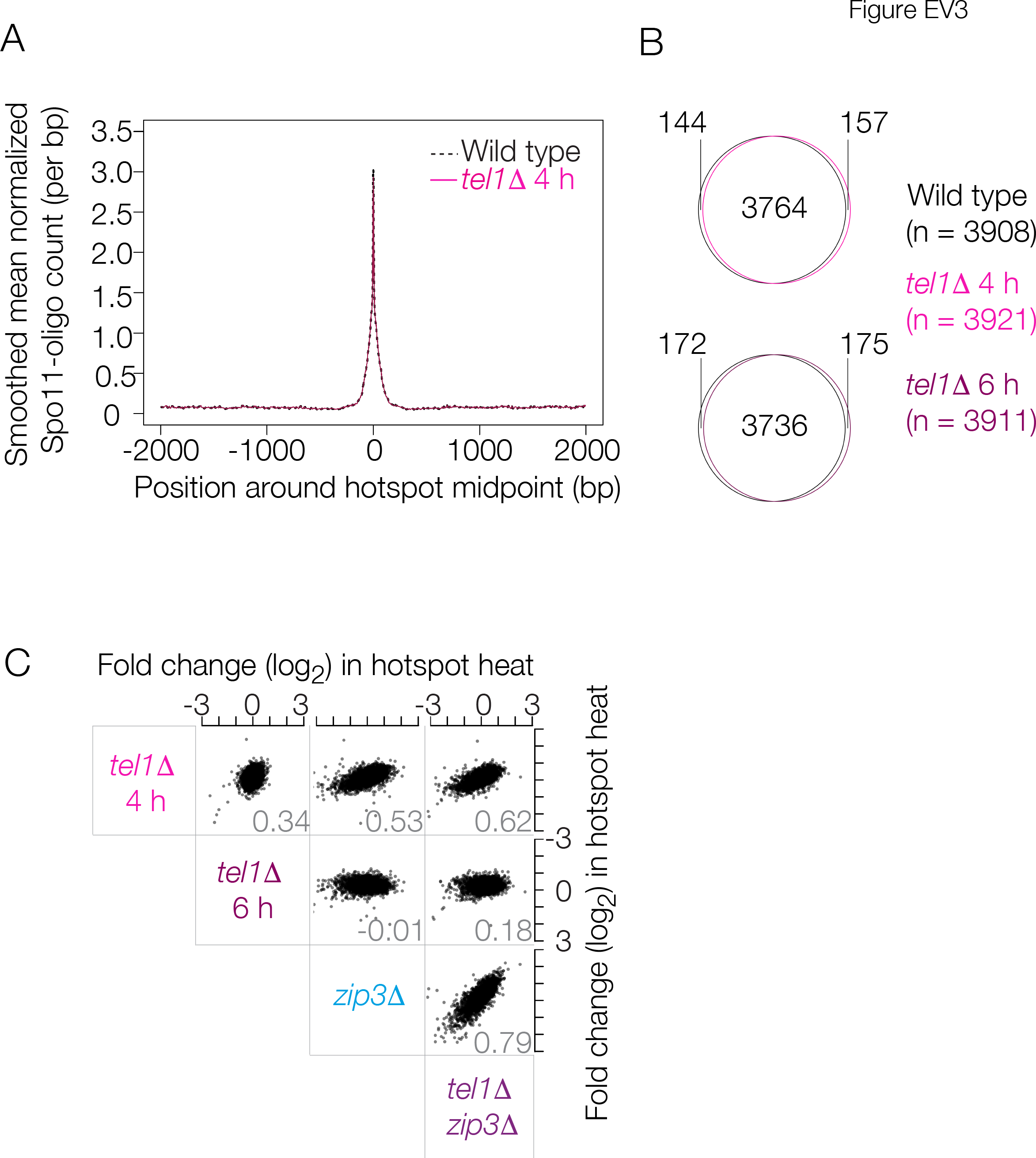
Hotspot-level effects of the *tellΔ* mutation. A: Hotspot widths are unaltered in *tel1Δ*. Spo11-oligos (normalized to RPM) were smoothed with a 201-bp Hann window and averaged around 3908 hotpots as defined in an averge Spo11-oligo map compiled from multiple wild-type datasets (see Methods). Identical results were obtained with the *tel1Δ* 6 h dataset. B: Venn diagrams depicting the high degree of overlap between hotspots called in wild-type and *tel1Δ* Spo11-oligo maps. C: Comparison between mutants for changes in activity of individual hotspots. Each point displays the pairwise comparison between mutant datasets for the fold change (log_2_, relative to wild type) in Spo11-oligo counts within a hotspot. Correlation coefficients (Pearson’s r) are shown at lower right of each plot.

**Fig. EV4.**
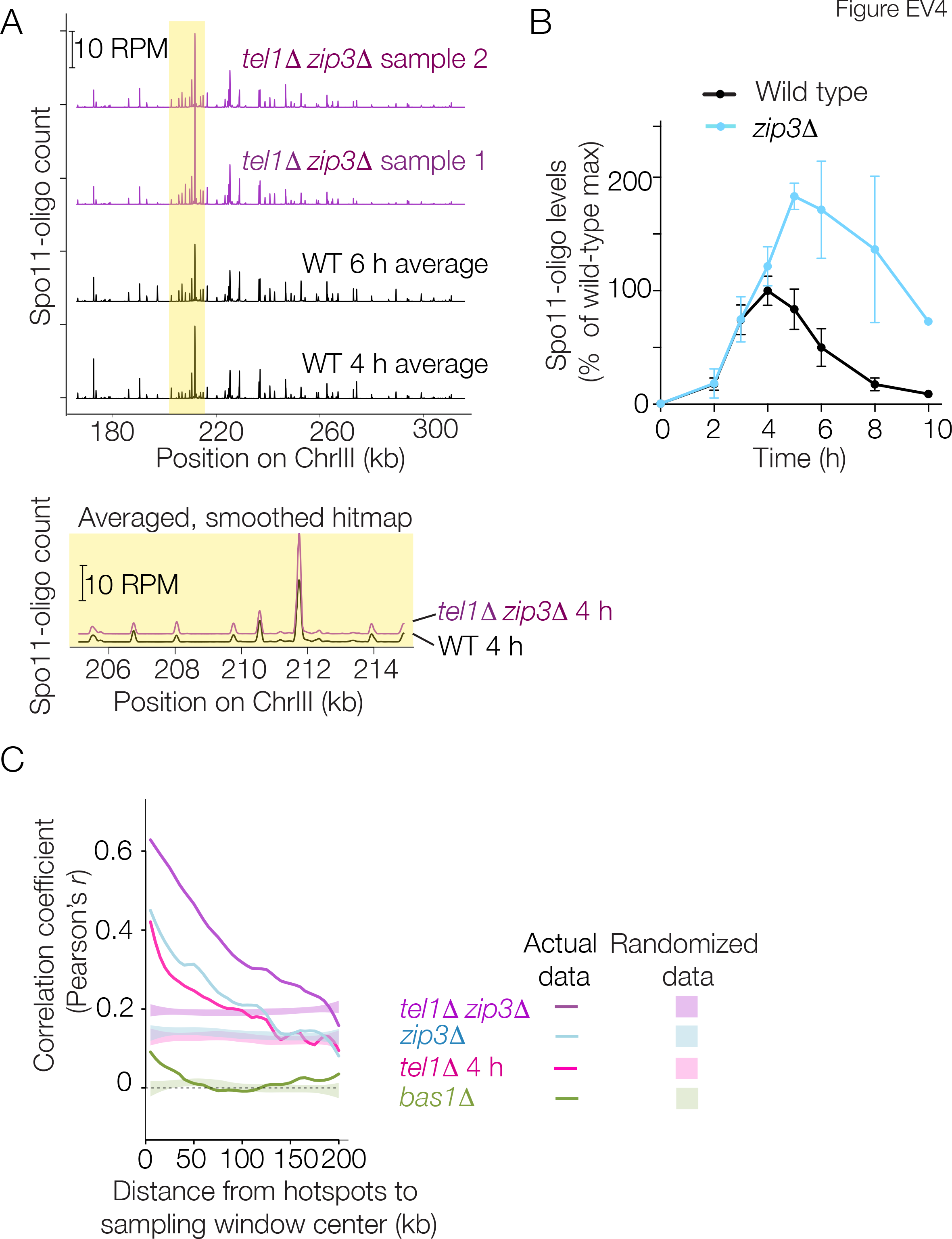
Spo11-oligo landscape in the *tellΔ zip3Δ* double mutant. A: Spo11-oligo map in *tel1Δ zip3Δ* mutant as displayed in **Fig. 4A**. B: Spo11-oligo quantification from (Thacker et al, 2014). C: Local domains of correlated behavior. The data from **Fig. 5B** are reproduced along with 95% confidence intervals for data randomized within-chromosome. For clarity, only the local regression lines fitted to the real data and not the individual points are shown.

**Fig. EV5.**
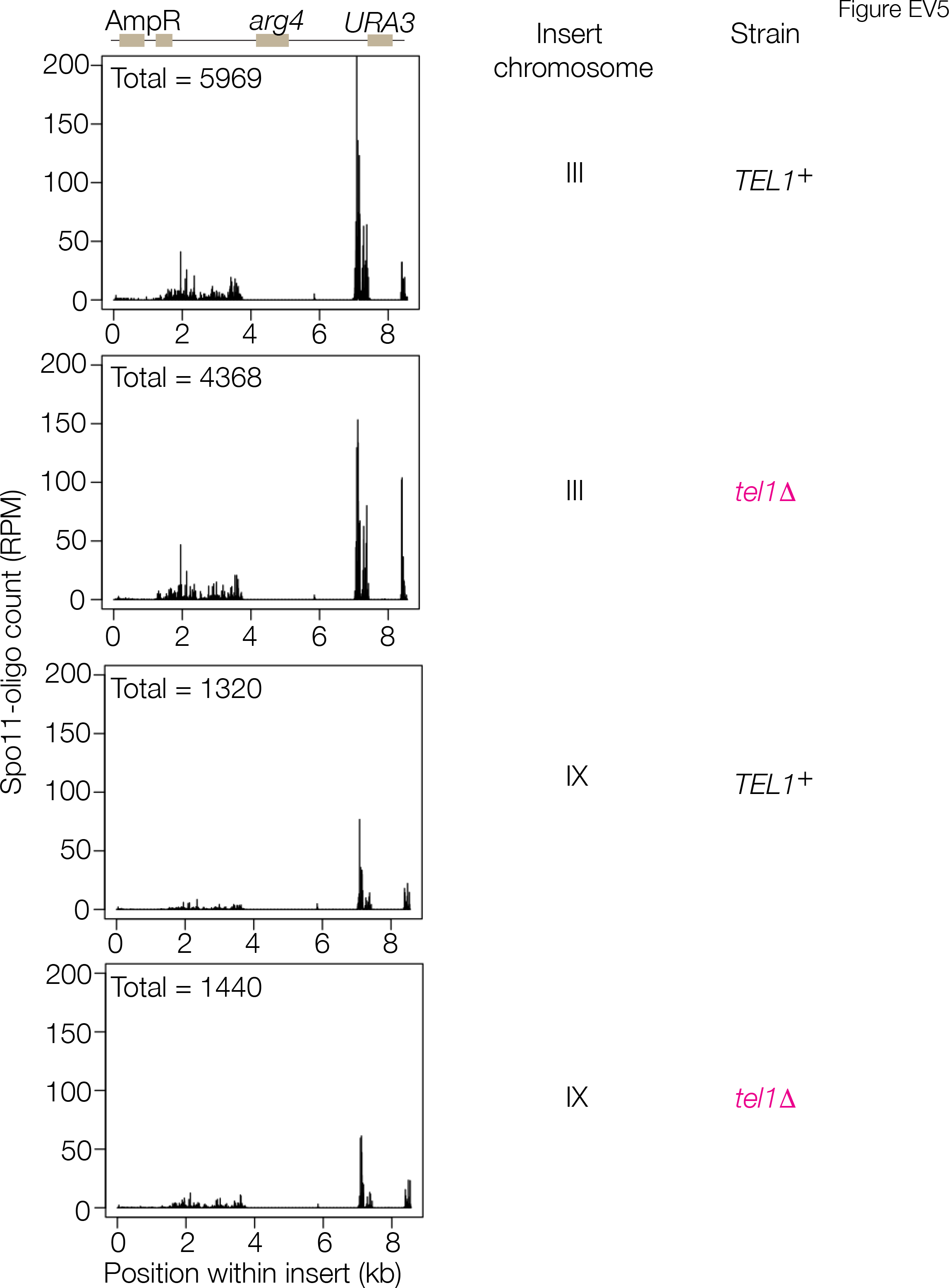
Uniquely mapping hits on the artificial hotspot insert. Top: Schematic of the artificial hotspot insert bearing mutant *arg4* and wild-type *URA3* genes within plasmid sequences (Wu & Lichten, 1995). Normalized unique Spo11-oligo maps within the insert are shown for the indicated *TEL1* genotypes and insert positions. Total normalized Spo11-oligo counts are indicated at upper left of each plot. Note that DSBs form principally within the plasmid backbone, as previously described (Wu & Lichten, 1995).

**Fig. EV6.**
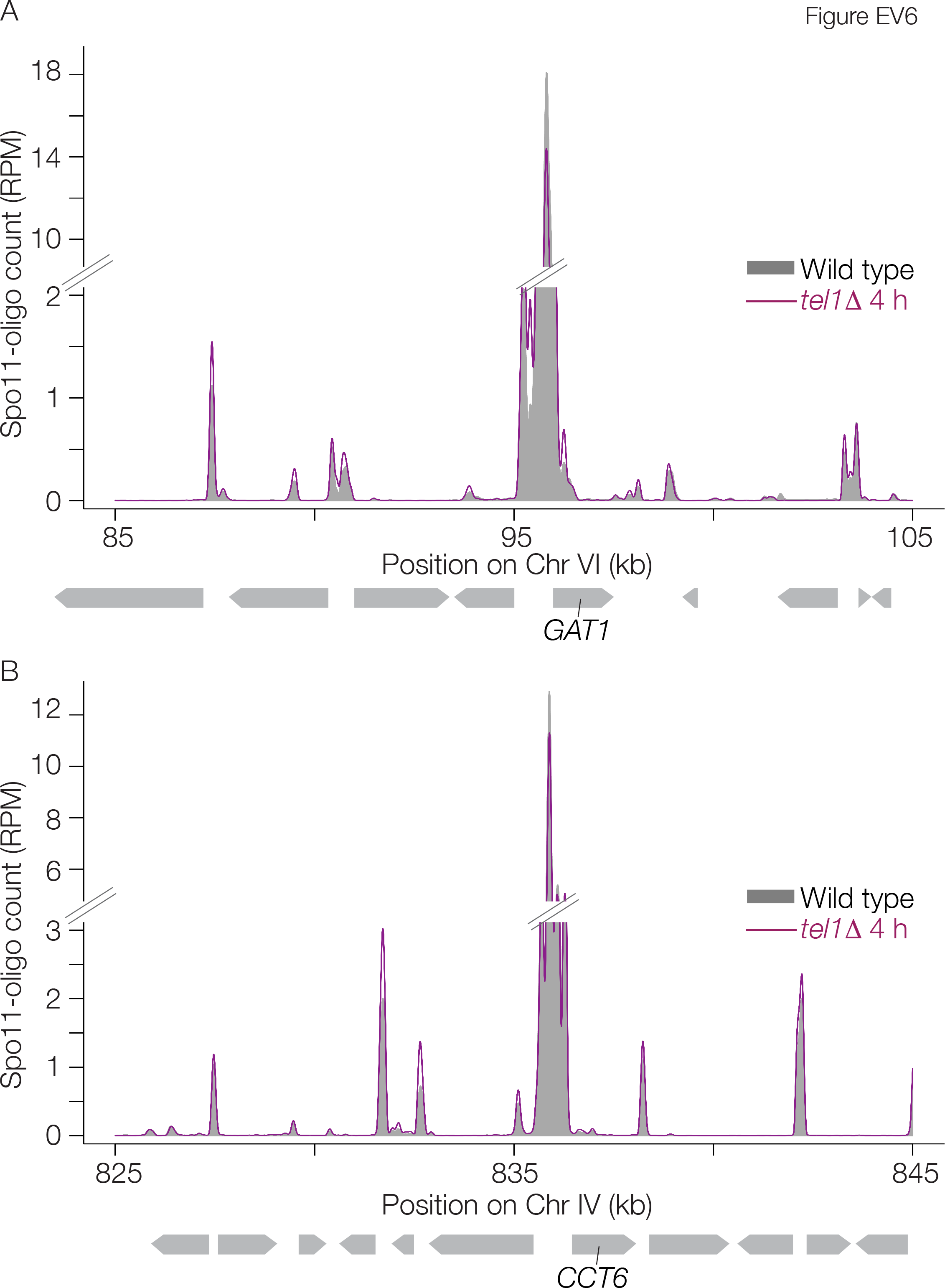
Hotspot competition around natural DSB hotspots. A,B: Spo11-oligo profiles around the *GAT1* and *CCT6* hotspots. Data were smoothed with a 201-bp Hann sliding window. Data are plotted as RPM without scaling to account for the overall increase in Spo11-oligo levels in *tel1Δ;* this allows the comparison between *tel1Δ* and wild type to provide a direct visualization of changes in hotspot activity relative to the genome-wide average.

**Table S1:**
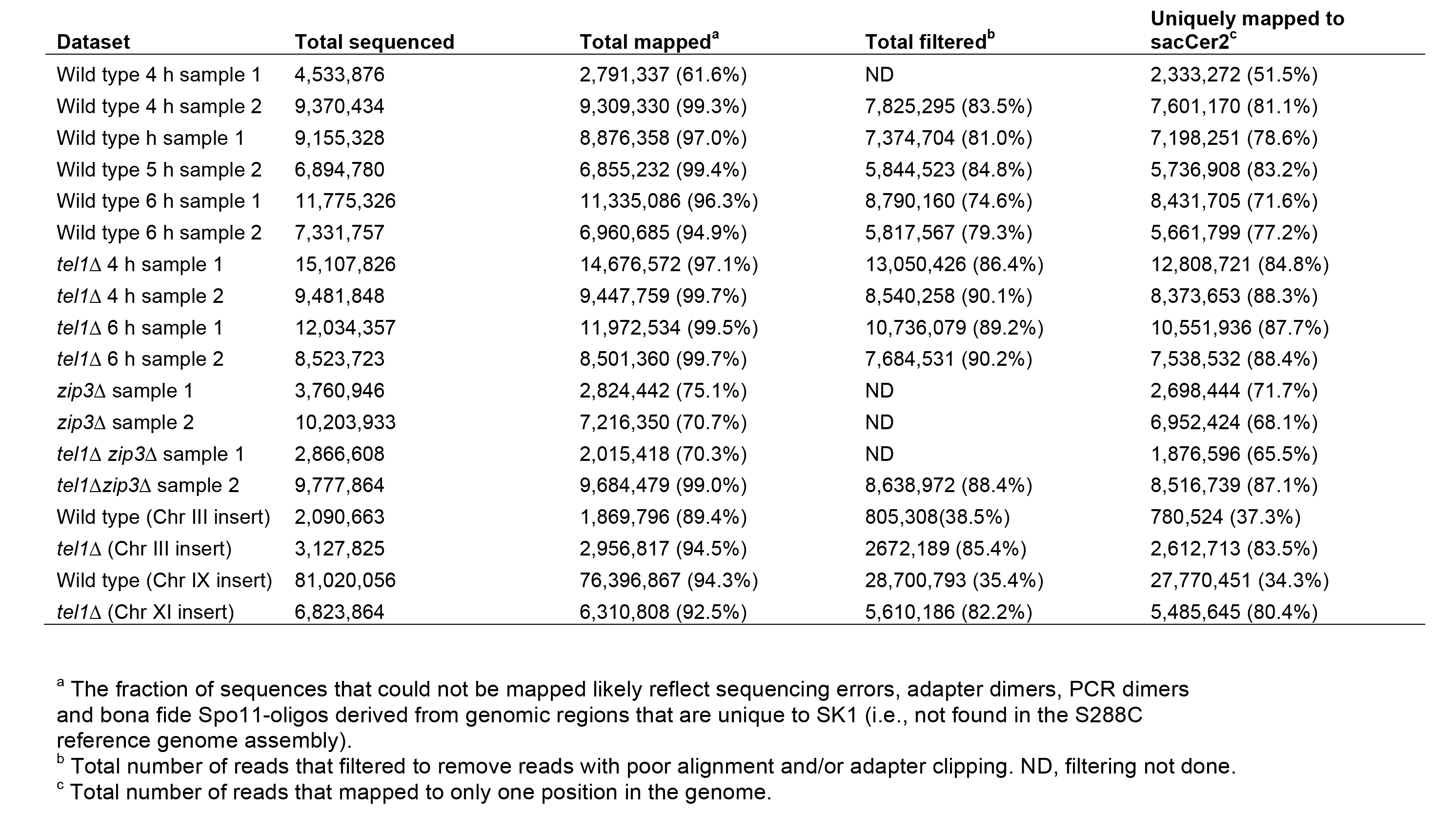
**Spo11-oligo mapping statistics.**

**Table S2:**
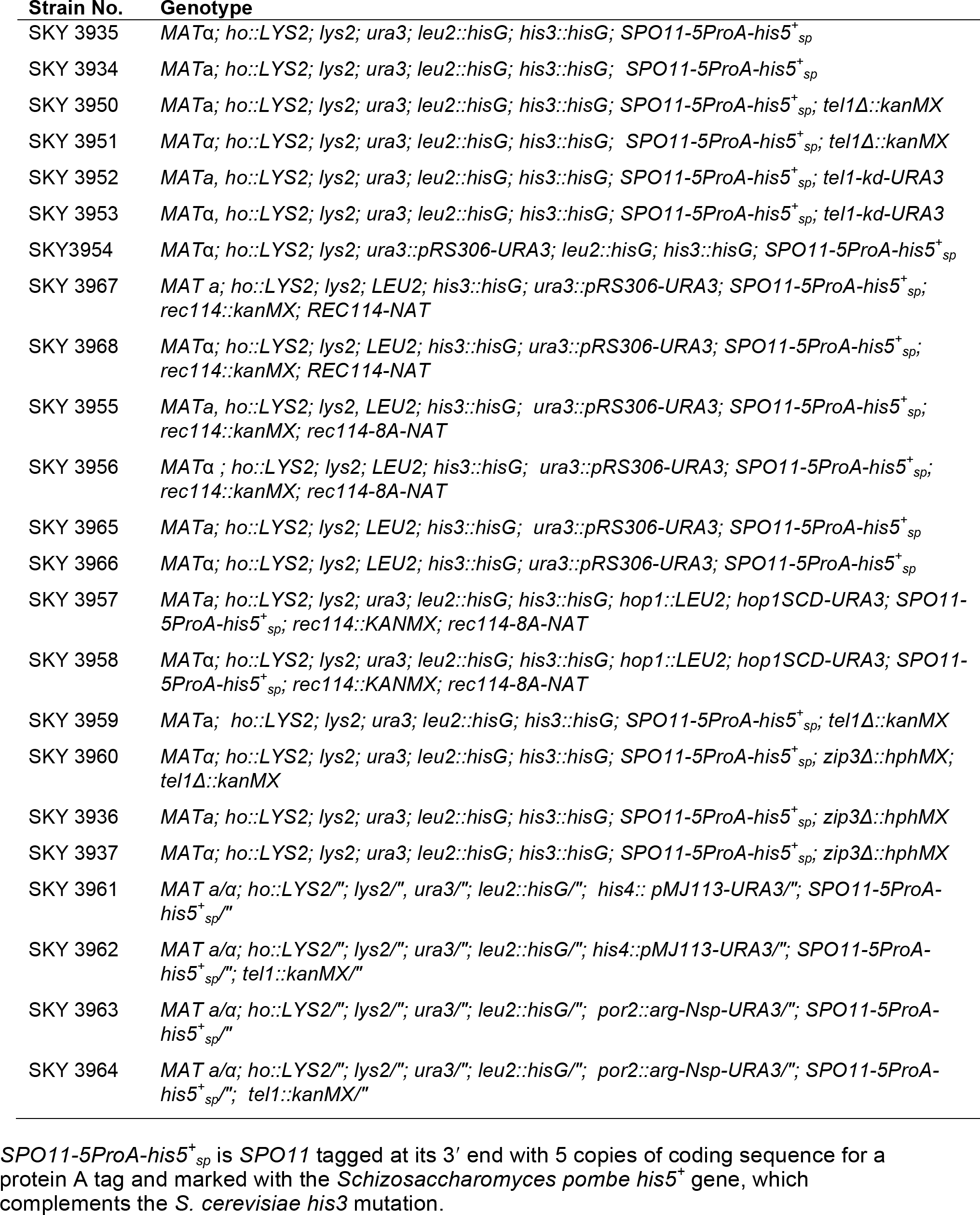
**Yeast strains**

## Numerical and Spatial Patterning of Yeast Meiotic DNA Breaks by Tell

Neeman Mohibullah and Scott Keeney

## Appendix Supplementary Methods

### Yeast Strains

Strains used in this study are from the SK1 background and are listed in **Table S2**. Tagging of Spo11 with 5 copies of a fragment of protein A was described previously and shown to be phenotypically normal in terms of DSB numbers (Thacker et al, 2014). The *tel1-kd* allele of Ma and Greider (Ma & Greider, 2009) (*tel1-D2612A,N2617A,D2631A*) was generated via QuickChange mutagenesis on a plasmid-borne 3′ fragment of *TEL1* which was subsequently linearized by digesting with *Nhel* and transformed into SKY 3934 and SKY 3935. Transformants were verified by preparing genomic DNA, digesting with *Clal*, separating on an agarose gel and Southern blotting using a probe to detect the *AVT5* open reading frame. Candidates with the correctly sized insertion were further verified by sequencing to ensure that the kinase-dead mutations had been retained. We recreated the *rec114-8A* allele described by (Carballo et al, 2013) (*S148A, T175A, T179A, S187A, S229A, T238A, S256A, S307A*) using synthetic DNA fragments spanning the mutated region of *REC114*. The fragments were stitched together using PCR sewing and used to replace the wild-type sequence on a plasmid-borne copy of *REC114* under control of the endogenous promoter (SacII to SpeI fragment around the *REC114* locus cloned into pAG25 (NAT)). The resulting mutant plasmid was cut with *MfeI* (within the *REC114* promoter) and transformed into *rec114A* cells lacking the *REC114* open reading frame. Transformants were verified by digesting genomic DNA with *XhoI* and *BlpI,* running the digested DNA on an agarose gel and screening for the correct size of insertion by probing a Southern of the digested DNA with probe to the *RRB1* ORF. The artificial hotspot was created as follows: fragments of the yeast genome containing coding regions from *HIS4* (+726 to +2062 with respect to ATG) or *POR2* (-380 to +1001 with respect to ATG) were cloned into the *EcoRI* sites of pMJ113 (Wu & Lichten, 1995), cut with either *XbaI* (*HIS4* hotspot) or *BstXI* (*POR2* hotspot) and transformed into the appropriate SK1 strains. The *hop1-scd* allele is *hop1-S298A, S311A, T318A* (Carballo et al, 2008) was provided by R. Cha.

### Time-course analysis of Spo11-oligo complexes

Cells were cultured and harvested from meiotic time courses as previously described (Thacker et al, 2014). Denaturing extracts were prepared from yeast cells using 10% trichloroacetic acid and agitation in a bead beater (three times for 30 s with cooling of tubes in ice-water between each cycle) using 600 μl of zirconium beads (BioSpec). Spo11 was immunoprecipitated via a C-terminal 5× Protein A tag. For the immunoprecipitation (IP) reactions we utilized magnetic beads conjugated to rabbit IgG prepared according to manufacturer’s instructions. 40 mg M-270 Epoxy Dynabeads (Thermo Fisher Scientific) were rotated overnight at 30°C with 400 μg rabbit IgG (Sigma cat. no. I5006) in the presence of 1 M ammonium sulphate in a final volume of 1.25 ml. The beads were subsequently washed with 1 ml of the following reagents in this order: 0.1 M sodium phosphate pH 7.4 (twice), 0.1 M glycine pH 2.5, 10 mM Tris-HCl pH 8.8, 100 mM freshly prepared triethylamine, 1× PBS (four times). The beads were resuspended in 1.25 ml of 1× PBS and 10 μl were used for each IP along with 200 μl of cell extract, rotated at 4°C overnight. End– labeling of Spo11-oligo complexes was carried out as previously described (Thacker et al, 2014) using 3-10 μCi of [*α*-^32^P] dCTP and terminal deoxynucleotidyl transferase (TdT). Spo11-oligo complexes were eluted by adding 25 μl of NUPAGE loading buffer (Thermo-Fisher) (diluted to 2× and supplemented with 100 mM dithiothreitol) and boiling for 5 min. End-labeled Spo11-oligo complexes were separated on a Novex 4-12% gradient denaturing polyacrylamide gel (Thermo-Fisher) then transferred onto PVDF membrane using the iBlot protocol (ThermoFisher) and visualized by phosphorimager. Western blotting to monitor free Spo11 was done using an anti-Protein A antibody (Sigma cat. no P2921), and chemiluminescent detection (ECL Prime, GE Life Sciences).

### Sizing of Spoil oligos

Spo11-oligo complexes were immunoprecipitated and labeled as above, the beads were washed with 1 ml IP buffer (2% Triton X-100, 30 mMTris-HCl, pH 8.0, 300 mM NaCl, 2 mM EDTA) and then with 1 ml Proteinase K buffer (100 mMTris-HCl, pH 7.4, 1 mM EDTA, 0.5% SDS, 1 mM CaCl_2_) lacking SDS. The beads were then transferred to 200 μl Proteinase K buffer and incubated with 1 μg Proteinase K and rotated overnight at 50°C. The supernatant was then transferred to a fresh tube with an equal volume of Proteinase K buffer lacking SDS and CaCl_2_ and ethanol precipitated. After removal of ethanol and washing with 70% ethanol, the precipitate was air-dried for 10 min, resuspended in formamide dye and run on a 15% denaturing polyacrylamide sequencing gel with radiolabeled 10-nt DNA ladder as a marker (Life Technologies).

## References

Anderson CM, Oke A, Yam P, Zhuge T, Fung JC (2015) Reduced Crossover Interference and Increased ZMM-Independent Recombination in the Absence of Tel1/ATM. PLoS Genet 11: e1005478

Argunhan B, Farmer S, Leung WK, Terentyev Y, Humphryes N, Tsubouchi T, Toyoizumi H, Tsubouchi H (2013) Direct and indirect control of the initiation of meiotic recombination by DNA damage checkpoint mechanisms in budding yeast. PLoS One 8: e65875

Blitzblau HG, Hochwagen A (2013) ATR/Mec1 prevents lethal meiotic recombination initiation on partially replicated chromosomes in budding yeast. Elife 2: e00844

Borde V, Wu T-C, Lichten M (1999) Use of a recombination reporter insert to define meiotic recombination domains on Chromosome III of Saccharomyces cerevisiae. Mol Cell Biol 19: 4832–4842

Bullard SA, Kim S, Galbraith AM, Malone RE (1996) Double strand breaks at the HIS2 recombination hot spot in Saccharomyces cerevisiae. Proc Natl Acad Sci U S A 93: 1305413059

Cannavo E, Cejka P (2014) Sae2 promotes dsDNA endonuclease activity within Mre11-Rad50-Xrs2 to resect DnA breaks. Nature 514: 122–125

Carballo JA, Johnson AL, Sedgwick SG, Cha RS (2008) Phosphorylation of the axial element protein Hop1 by Mec1/Tel1 ensures meiotic interhomolog recombination. Cell 132: 758–770

Carballo JA, Panizza S, Serrentino ME, Johnson AL, Geymonat M, Borde V, Klein F, Cha RS (2013) Budding yeast ATM/ATR control meiotic double-strand break (DSB) levels by down-regulating Rec114, an essential component of the DSB-machinery. PLoS Genet 9: e1003545

Cartagena-Lirola H, Guerini I, Viscardi V, Lucchini G, Longhese MP (2006) Budding Yeast Sae2 is an In Vivo Target of the Mec1 and Tel1 Checkpoint Kinases During Meiosis. Cell Cycle 5: 1549–1559

Cooper TJ, Garcia V, Neale MJ (2016) Meiotic DSB patterning: A multifaceted process. Cell Cycle 15: 13–21

Daniel JA, Pellegrini M, Lee BS, Guo Z, Filsuf D, Belkina NV, You Z, Paull TT, Sleckman BP, Feigenbaum L, Nussenzweig A (2012) Loss of ATM kinase activity leads to embryonic lethality in mice. J Cell Biol 198: 295–304

Fan QQ, Xu F, White MA, Petes TD (1997) Competition between adjacent meiotic recombination hotspots in the yeast Saccharomyces cerevisiae. Genetics 145: 661–670

Fukuda T, Kugou K, Sasanuma H, Shibata T, Ohta K (2008) Targeted induction of meiotic double-strand breaks reveals chromosomal domain-dependent regulation of Spo11 and interactions among potential sites of meiotic recombination. Nucleic Acids Res 36: 984–997

Garcia V, Gray S, Allison RM, Cooper TJ, Neale MJ (2015) Tel1(ATM)-mediated interference suppresses clustered meiotic double-strand-break formation. Nature 520: 114–118

Garcia V, Phelps SE, Gray S, Neale MJ (2011) Bidirectional resection of DNA double-strand breaks by Mre11 and Exo1. Nature 479: 241–244

Hayashi M, Mlynarczyk-Evans S, Villeneuve AM (2010) The synaptonemal complex shapes the crossover landscape through cooperative assembly, crossover promotion and crossover inhibition during Caenorhabditis elegans meiosis. Genetic 186: 45–58

Hunter N, Kleckner N (2001) The single-end invasion: an asymmetric intermediate at the double-strand break to double-holliday junction transition of meiotic recombination. Cell 106: 59–70

Joshi N, Brown MS, Bishop DK, Borner GV (2015) Gradual implementation of the meiotic recombination program via checkpoint pathways controlled by global DSB levels. Mol Cell 57: 797–811

Joyce EF, Pedersen M, Tiong S, White-Brown SK, Paul A, Campbell SD, McKim KS (2011) Drosophila ATM and ATR have distinct activities in the regulation of meiotic DNA damage and repair. J Cell Biol 195: 359–367

Kauppi L, Barchi M, Lange J, Baudat F, Jasin M, Keeney S (2013) Numerical constraints and feedback control of double-strand breaks in mouse meiosis.Genes Dev 27: 873–886

Keeney S, Lange J, Mohibullah N (2014) Self-organization of meiotic recombination initiation: general principles and molecular pathways. Annu Rev Genet 48: 187–214

Klein F, Mahr P, Galova M, Buonomo SB, Michaelis C, Nairz K, Nasmyth K (1999) A central role for cohesins in sister chromatid cohesion, formation of axial elements, and recombination during yeast meiosis. Cell 98: 91–103

Kugou K, Fukuda T, Yamada S, Ito M, Sasanuma H, Mori S, Katou Y, Itoh T, Matsumoto K, Shibata T, Shirahige K, Ohta K (2009) Rec8 guides canonical Spo11 distribution along yeast meiotic chromosomes. Mol Biol Cell 20: 3064–3076

Lam I, Keeney S (2015a) Mechanism and regulation of meiotic recombination initiation. Cold Spring Harb Perspect Biol 7: a016634

Lam I, Keeney S (2015b) Nonparadoxical evolutionary stability of the recombination initiation landscape in yeast. Science 350: 932–937

Lange J, Pan J, Cole F, Thelen MP, Jasin M, Keeney S (2011) ATM controls meiotic double-strand-break formation. Nature 479: 237–240

Lynn A, Soucek R, Borner GV (2007) ZMM proteins during meiosis: crossover artists at work. Chromosome Res 15: 591–605

Ma Y, Greider CW (2009) Kinase-independent functions of TEL1 in telomere maintenance. Mol Cell Biol 29: 5193–5202

Neale MJ, Pan J, Keeney S (2005) Endonucleolytic processing of covalent protein-linked DNA double-strand breaks. Nature 436: 1053–1057

Pan J, Sasaki M, Kniewel R, Murakami H, Blitzblau HG, Tischfield SE, Zhu X, Neale MJ, Jasin M, Socci ND, Hochwagen A, Keeney S (2011) A hierarchical combination of factors shapes the genome-wide topography of yeast meiotic recombination initiation. Cell 144: 719–731

Robine N, Uematsu N, Amiot F, Gidrol X, Barillot E, Nicolas A, Borde V (2007) Genome-wide redistribution of meiotic double-strand breaks in Saccharomyces cerevisiae. Mol Cell Biol 27: 1868–1880

Shiloh Y, Ziv Y (2013) The ATM protein kinase: regulating the cellular response to genotoxic stress, and more. Nat Rev Mol Cell Biol 14: 197–210

Terasawa M, Ogawa T, Tsukamoto Y, Ogawa H (2008) Sae2p phosphorylation is crucial for cooperation with Mre11p for resection of DNA double-strand break ends during meiotic recombination in Saccharomyces cerevisiae. Genes Genet Syst 83: 209–217

Thacker D, Mohibullah N, Zhu X, Keeney S (2014) Homologue engagement controls meiotic DNA break number and distribution. Nature 510: 241–246

Wojtasz L, Daniel K, Roig I, Bolcun-Filas E, Xu H, Boonsanay V, Eckmann CR, Cooke HJ, Jasin M, Keeney S, McKay MJ, Toth A (2009) Mouse HORMAD1 and HORMAD2, two conserved meiotic chromosomal proteins, are depleted from synapsed chromosome axes with the help of TRIP13 AAA-ATPase. PLoS Genet 5: e1000702

Wu T-C, Lichten M (1995) Factors that affect the location and frequency of meiosis-induced double-strand breaks in Saccharomyces cerevisiae. Genetics 140: 55–66

Xu L, Kleckner N (1995) Sequence non-specific double-strand breaks and interhomolog interactions prior to double-strand break formation at a meiotic recombination hot spot in yeast. EMBO J 14: 5115–5128

Yamamoto K, Wang Y, Jiang W, Liu X, Dubois RL, Lin CS, Ludwig T, Bakkenist CJ, Zha S (2012) Kinase-dead ATM protein causes genomic instability and early embryonic lethality in mice. J Cell Biol 198: 305–313

Zhang L, Kim KP, Kleckner NE, Storlazzi A (2011) Meiotic double-strand breaks occur once per pair of (sister) chromatids and, via Mec1/ATR and Tel1/ATM, once per quartet of chromatids. Proc Natl Acad Sci U S A 108: 20036–20041

Zhu X, Keeney S (2015) High-Resolution Global Analysis of the Influences of Bas1 and Ino4 Transcription Factors on Meiotic DNA Break Distributions in Saccharomyces cerevisiae. Genetics 201: 525–542

